# Biosynthesis of the redox cofactor mycofactocin comprises oligoglycosylation by MftF in *Mycolicibacterium smegmatis*

**DOI:** 10.1101/821413

**Authors:** Luis Peña-Ortiz, Ana Patrícia Graça, Huijuan Guo, Daniel Braga, Tobias G. Köllner, Lars Regestein, Christine Beemelmanns, Gerald Lackner

**Affiliations:** Junior Research Group Synthetic Microbiology, Leibniz Institute for Natural Product Research and Infection Biology – Hans Knöll Institute, Beutenbergstr. 11a, 07745 Jena, Germany; Friedrich Schiller University, Jena, Germany; Junior Research Group Chemical Biology of Microbe-Host Interactions, Leibniz Institute for Natural Product Research and Infection Biology – Hans Knöll Institute, Beutenbergstr. 11a, 07745 Jena, Germany; Max Planck Institute for Chemical Ecology, Department of Biochemistry, Hans-Knöll-Str. 8, 07745 Jena, Germany; Bio Pilot Plant, Leibniz Institute for Natural Product Research and Infection Biology – Hans Knöll Institute, Beutenbergstr. 11a, 07745 Jena, Germany

**Keywords:** Mycofactocin, cofactor, *Mycobacterium*, metabolomics, mass spectrometry (MS), alcohol metabolism, drug target

## Abstract

Mycofactocin (MFT) is a redox cofactor involved in alcohol metabolism of mycobacteria including *Mycobacterium tuberculosis*. In recent years, a preliminary biosynthetic model of MFT has been established by *in-vitro* studies, while the final structure of MFT remained elusive. Here, we report the discovery of MFT by metabolomics and establish a model of its biosynthesis in *Mycolicibacterium smegmatis*. Structure elucidation revealed that MFT is decorated with up to nine β-1,4-linked glucose residues. Dissection of biosynthetic genes demonstrated that the oligoglycosylation is catalyzed by the glycosyltransferase MftF. Furthermore, we confirm the cofactor function of MFT by activity-based metabolic profiling using the carveol dehydrogenase LimC and show that the MFT pool expands during cultivation on ethanol. Our results close an important gap of knowledge, will guide future studies into the physiological roles of MFT in bacteria and may inspire its utilization as a biomarker or potential drug target to combat mycobacterial diseases.

## Introduction

Coenzymes are small molecules that are indispensable for the catalytic activity of many enzymes. While coenzymes like NAD^+^ or FAD are ubiquitous in nature and are essential for the core metabolism of all forms of life, specialized cofactors like pyrroloquinoline quinone (PQQ)^1^ and coenzyme F_420_^2^ are restricted to certain microbial phyla, but typically involved in extraordinary metabolic processes like methylotrophy, methanogenesis, or detoxification processes. Mycobacteria are particularly rich in unusual redox cofactors and antioxidants that contribute to redox balance and metabolic plasticity. For instance, mycothiol^3, 4^ or ergothioneine^5^ protect *Mycobacterium tuberculosis* from oxidative stress and support detoxification pathways while coenzyme F_420_ is involved in a response to nitrosative stress^6^. Pathogenic mycobacteria are exposed to a variety of carbon sources, reactive oxygen species, and reactive nitrogen species inside the host organism. Furthermore, persistent *M. tuberculosis* may enter a dormant state characterized by low metabolic activity and reduced susceptibility to antibiotics, where survival or death are governed by complex redox signaling pathways and alterations of electron transport processes. Moreover, some antimycobacterial drugs are administered as prodrugs and will only develop bioactivity upon transfomation, e.g. protomanid^7^, which is activated by a coenzyme F_420_-dependent reductase^8^. Hence, unusual cofactors that assist such biochemical processes are critical for survival, virulence, and drug resistance of *M. tuberculosis*^9^. The emergence of extensively drug-resistant tuberculosis (XDR-TB) is one of the major threats to human health worldwide and highlights the urgency of the exploration of metabolic systems that may influence the fitness of a pathogen during infection and modulate antibiotics activity.

Mycofactocin (MFT)^10^ is a putative redox-cofactor whose existence has been postulated on the basis of comparative genomics and bioinformatics^11^. Its final molecular identity and structure, however, have remained elusive to date. The MFT biosynthetic gene cluster is highly conserved and wide-spread among mycobacteria. The inactivation of the MFT gene locus in the model species *Mycolicibacterium smegmatis* (synonym: *Mycobacterium smegmatis*) as well as *M. tuberculosis* resulted in the inability of the mutants to utilize ethanol as a sole source of carbon and further disturbances of mycobacterial redox homeostasis were revealed^12^. Involvement of MFT in methanol metabolism was reported as well^13^. These recent results strongly support the hypothesis that MFT is a redox cofactor and might represent a fitness factor of mycobacteria during some stages of infection.

The architecture of the MFT gene cluster (Fig. 1a) suggested that the resulting natural product is a ribosomally synthesized and post-translationally modified peptide (RiPP)^14^. Several *in-vitro* studies have contributed to a biosynthetic model of MFT (Fig. 1b): The precursor peptide MftA consisting of 32 amino acids is produced by the ribosome and bound by its chaperone MftB. Subsequently, the terminal core peptide consisting of Val and Tyr is oxidatively decarboxylated and cyclized by the radical SAM enzyme MftC^15, 16, 17^. The resulting cyclic core structure is released by the peptidase MftE^18^ forming 3-amino-5-[(*p-*hydroxyphenyl)methyl]-4,4-dimethyl-2-pyrrolidinone (AHDP)^19^. Just recently, it was shown that MftD, an enzyme homologous to the L-lactate dehydrogenase LldD2^20^, catalyzes the oxidative deamination of AHDP to yield pre-mycofactocin (PMFT)^21^. The same study demonstrated by voltammetry that the α-keto amide moiety of PMFT is redox-active and can be reduced to PMFTH_2_ (midpoint potential: −255 mV). Efficient reduction was also achieved by the action of carveol dehydrogenase using carveol as an electron donor *in vitro*^21^. Therefore, PMFT likely represents the redox-center of MFT, as riboflavin is the redox-active core of FMN and FAD.

**Figure 1.**
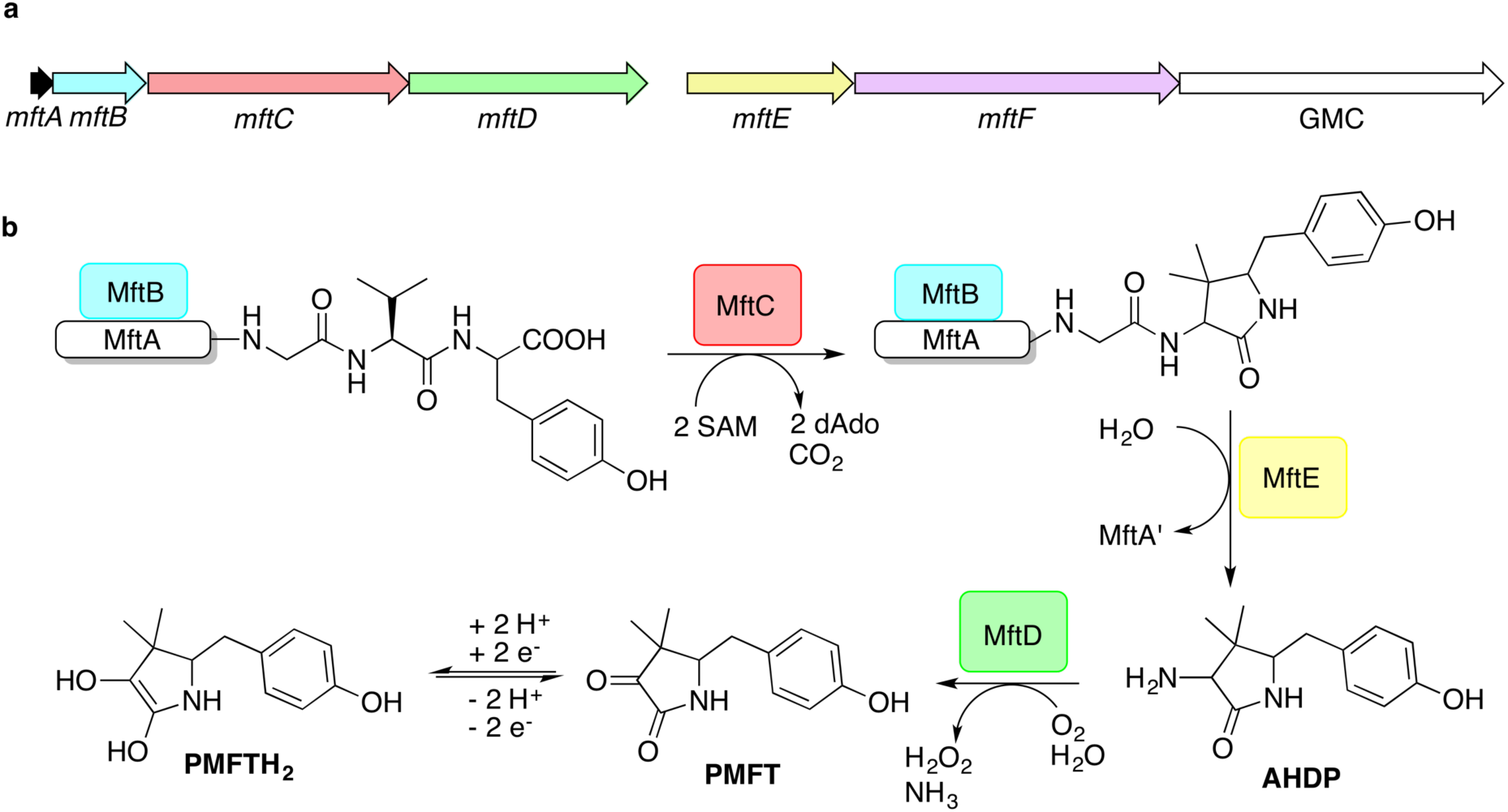
Biosynthesis of mycofactocin. **a** Schematic representation of the MFT biosynthetic gene cluster of *M. smegmatis*. Arrows present coding regions of genes *mftA-F*. GMC: glucose-methanol-choline oxidoreductase. **b** Current biosynthesis model of MFT revealed by *in-vitro* studies. The precursor peptide MftA is bound by its chaperone MftB. The rSAM enzyme MftC catalyzes oxidative decarboxylation and cyclization of the core peptide consisting of a C-terminal Val-Tyr dipeptide. The peptidase MftE releases the core to form 3-amino-5-[(*p*-hydroxyphenyl)methyl]-4,4-dimethyl-2-pyrrolidinone (AHDP). MftD performs oxidative deamination of AHDP yielding pre-mycofactocin (PMFT), the presumed redox-active core. (P)MFT is reduced to (P)MFTH_2_ (mycofactocinol) by unknown oxidoreductases. dAdo: 5’-deoxyadenosine, SAM: *S*-adenosyl methionine.

Although these current hypotheses are plausible, all of these known metabolic intermediates have just been observed *in vitro* and could therefore represent artifacts. The verification of their relevance *in vivo* was urgently desired and the final steps of MFT biosynthesis as well as the function of the *mftF* gene awaited experimental elucidation. In this study, we confirm the current biosynthetic model of MFT *in vivo* and detected several novel oligoglycosylated MFT congeners. We also show that the sugar chain is a β-1,4-glucan and provide first evidence that glycosylation is performed by the glycosyltransferase MftF. Finally, we show dependence of MFT formation on ethanol and corroborate its cofactor function by activity-based metabolic profiling.

## Results and Discussion

### Discovery of mycofactocins by metabolomics

In order to identify potential mycofactocin congeners in mycobacteria, we used the fast-growing and weakly pathogenic species *M. smegmatis* MC^2^ 155 as a model organism and developed a metabolomics approach combining metabolic induction and labeling to specifically trace MFT congeners. Assuming that MFT production would be stimulated by alcohols, we cultivated bacteria in media containing 10 g L^−1^ ethanol. Furthermore, we used stable isotope labeling to obtain candidate molecules compatible with the proposed biosynthetic pathway: Since the C-terminal core peptide of MftA is composed of Val and Tyr, we reasoned that MFT congeners could be specifically labeled by feeding L-Val-^13^C_5_ and L-Tyr-^13^C_9_. Intracellular contents were extracted and analyzed by liquid chromatography coupled with high-resolution mass spectrometry (LC-MS). Compounds were detected by *in-silico* grouping of co-eluting isotopic peaks and adducts (feature finding). Afterwards, ^13^C-labeled compounds were deduced computationally (Supplementary Table 1). According to the established biosynthetic pathway we expected 13 carbons to remain ^13^C-labeled after oxidative decarboxylation of the Val-Tyr core peptide. We therefore searched for compounds that displayed an exchange of exactly 13 carbons, resulting in a mass shift of +13.04362 Da (Supplementary Fig. 1). After removal of low-quality hits, this approach revealed a list of only twelve candidate compounds. Strikingly, the exact mass and proposed sum formula of three of these compounds corresponded to known intermediates of MFT, namely AHDP, PMFT as well as PMFTH_2_, indicating that our strategy was viable. In addition to these compounds, several labeled molecules with increasing molecular weight were detected. Some co-eluting candidates with a mass difference of +17.02654 could be explained as NH_4_^+^ adducts of each other. The remaining nine candidate compounds (Table 1) were grouped based on their chromatographic retention times, eluting closely to either PMFT or PMFTH_2_ (approx. 7.2 min and 6.9 min, respectively). Intriguingly, members of the two groups could be arranged in pairs with a mass difference of two hydrogen atoms leading us to assume that each group represented derivatives of either PMFT or PMFTH_2_. Thus, we termed these two clusters of molecules mycofactocinones (MFT) and mycofactocinols (MFTH_2_), respectively. Notably, some mycofactocinols eluted as two chromatographically separated isomers. For instance, the dominant PMFTH_2_ peak eluted at 6.8 min, while the minor isomer eluted at 6.5 min. These two compounds displayed highly similar MS/MS spectra (Supplementary Fig. 2) and most likely represent tautomeric forms. For the sake of simplicity, only the more prevalent isomer was considered during metabolomics studies.

**Table 1:** Curated list of MFT candidate molecules obtained by stable isotope labeling after induction with 10 g L^−1^ ethanol. Mycofactocinols (MFT-nH_2_) co-elute with PMFTH_2_ and represent (oligo-)glycosylated forms of PMFTH_2_. Mycofactocinons (MFT-n) co-elute with PMFT and represent (oligo-)glycosylated forms of PMFT. MMFT-n: methylmycofactocinones, MMFT-nH_2_: methylmycofactocinones. n represents the number of sugar moieties. Compounds revealed by stable isotope labeling (bold) were determined by Compound Discoverer 3.0 after feeding of *M. smegmatis* with L-Val-^13^C_9_ and L-Tyr-^13^C_9_. Values represent the mean of 4 biological replicates. All labeled compounds are shown in Supplementary Table 1, all MFT congeners revealed in this study are shown in Supplementary Table 2. RT: retention time.

| Name | Sum formula | Exact mass (measured) | RT [min] | Area (mean) |
| --- | --- | --- | --- | --- |
| **Precursors** |  |  |  |  |
| AHDP | C<sub>13</sub> H<sub>18</sub> N<sub>2</sub> O<sub>2</sub> | 234.13683 | 6.53 | 44862 |
| PMFTH<sub>2</sub> | C<sub>13</sub> H<sub>17</sub> N O<sub>3</sub> | 235.12084 | 6.88 | 49905 |
| PMFT | C<sub>13</sub> H<sub>15</sub> N O<sub>3</sub> | 233.10519 | 7.22 | 100832 |
| **Mycofactocinols (MFT-nH<sub>2</sub>)** |  |  |  |  |
| MFT-1H<sub>2</sub> | C<sub>19</sub> H<sub>27</sub> N O<sub>8</sub> | 397.17367 | 6.88 | 53413 |
| **Methylmycofactocinols (MMFT-nH<sub>2</sub>)** |  |  |  |  |
| MMFT-2H<sub>2</sub> | C<sub>26</sub> H<sub>39</sub> N O<sub>13</sub> | 573.24214 | 6.89 | 5722 |
| MMFT-7H<sub>2</sub> | C<sub>56</sub> H<sub>89</sub> N O<sub>38</sub> | 1383.50626 | 6.89 | 340064 |
| MMFT-8H<sub>2</sub> | C<sub>62</sub> H<sub>99</sub> N O<sub>43</sub> | 1545.55908 | 6.86 | 550390 |
| **Methylmycofactocinones (MMFT-n)** |  |  |  |  |
| MMFT-7 | C<sub>56</sub> H<sub>87</sub> N O<sub>38</sub> | 1381.49061 | 7.25 | 185407 |
| MMFT-8 | C<sub>62</sub> H<sub>97</sub> N O<sub>43</sub> | 1543.54343 | 7.21 | 335701 |

During further experiments, we performed MS/MS networking (Fig. 2), an approach that clusters compounds based on similarity of their MS/MS fragmentation pattern and therefore potentially related chemical scaffolds^22^. Interestingly, candidates retrieved from ^13^C-labeling experiments clustered with further putative MFT congeners. The mass difference between the first candidate mycofactocinol (MFT-1H_2_) with an exact mass of 397.17395 Da and PMFTH_2_ was +162.05303 Da, which corresponded to a hexose sugar. Furthermore, the MS/MS spectrum of MFT-1H_2_ (Fig. 2) showed a fragment ion that corresponded to the mass of the putative aglycon (*m/z* 236.13 [M+H]^+^), thus supporting the assumption that MFT-1H_2_ was a glycosylated derivative of PMFTH_2_. MS/MS networking also revealed a recurrent mass difference of 14.01565 between compounds, indicating that methylation might occur as well. We thus assumed that all MFT candidate molecules revealed by both ^13^C-labeling and MS/MS networking could be explained as (oligo-)glycosylated or glycosylated and monomethylated species of PMFT(H_2_). In analogy to coenzyme F_420_-n, where n indicates the number of glutamyl residues in the side chain^2^, we named the glycosylated molecules MFT-n(H_2_) with n representing the number of sugar moieties. Monomethylated species were termed methylmycofactocinones (MMFT-n) and methylmycofactocinols (MMFT-nH_2_), respectively. A targeted search for theoretical mass traces revealed additional members of the MFT-n(H_2_) and MMFT-n(H_2_) series (Supplementary Table 2).

**Figure 2.**
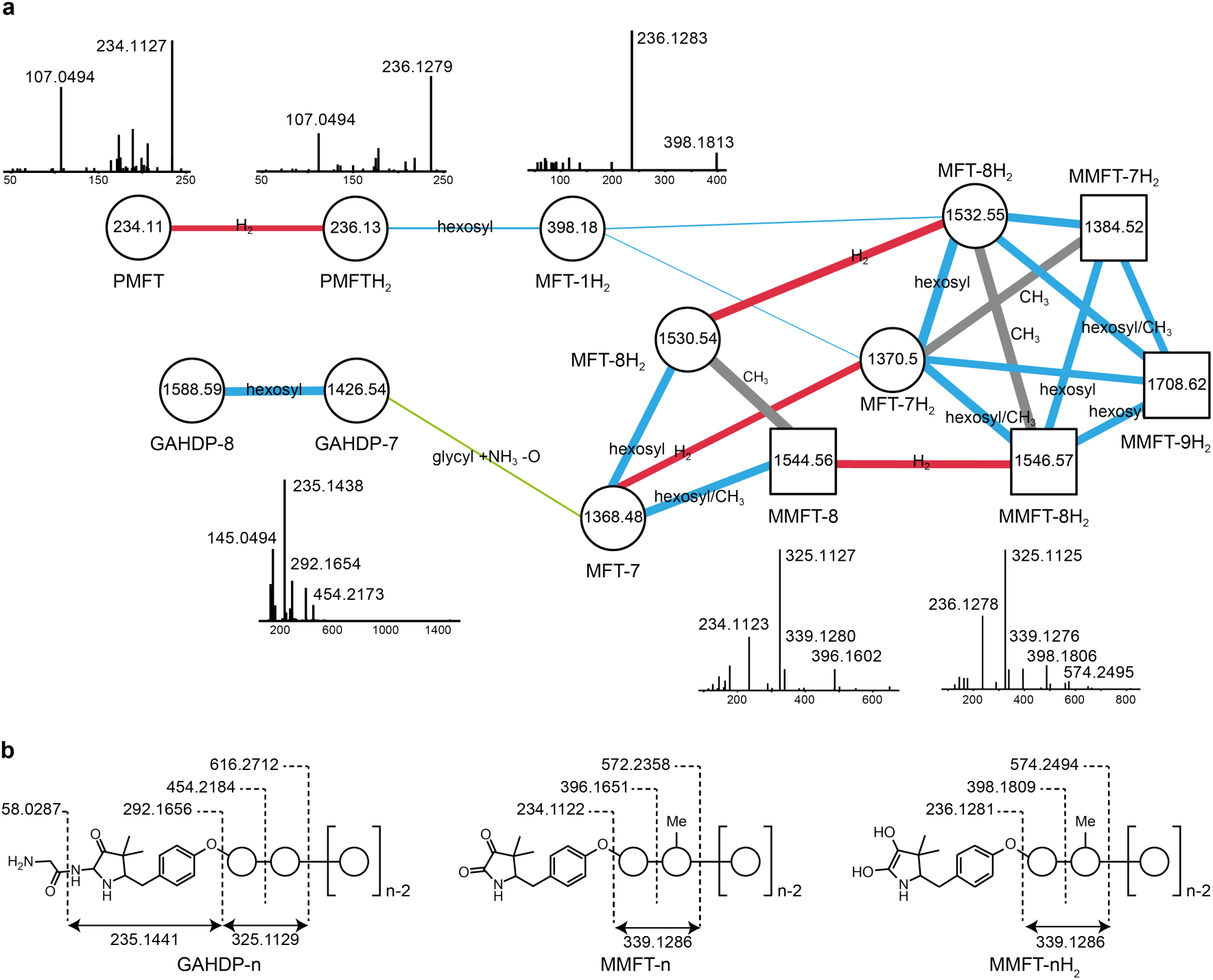
Discovery and tandem mass spectrometry of MFT congeners. **a** Molecular network of MFT congeners. Nodes (circles) represent chemical compounds. Internal node labels display the precursor mass of compounds (*m/z* [M+H^+^]). External node labels show proposed compound annotations. Edges represent relationships in terms of shared MS/MS fragments. Edge labels show proposed modifications based on precursor mass shifts (blue: hexosylation, red: oxidation/reduction, grey: methylation). Line widths of edges mirror cosine distances. Representative MS/MS spectra of corresponding precursor ions are shown above or below nodes. **b** Schematic representation of mass fragmentation patterns of GAHDP-n, MMFT-nH2 and MMFT-n. Numbers indicate mass-to-charge ratios (*m/z*) of fragments observed. Circles represent hexose moieties. Me: methyl group.

In accordance to the expectation, the corresponding mycofactocinones (e.g. MMFT-8) exhibited MS/MS fragments that showed a systematic shift by −2.0016 (e.g., *m/z* 234.11, 396.16, 572.24) (Fig. 2a) demonstrating that the reduction/oxidation indeed takes place in the PMFT moiety. Oligoglycosylation with up to n = 9 saccharide units was detected, while seven and eight units appeared to be the most dominant form in terms of area under the curve. Although we observed both methylated (MMFT) and unmethylated (MFT) sugar chains, the MMFT series appeared to be more prominent. Only monomethylated species were found. Furthermore, only MFT-1(H_2_), but not MMFT-1(H_2_) was detected, suggesting that the first sugar was not methylated. Mass fragmentation of MMFT-n(H_2_) species was well in agreement with the assumption that the second sugar was a hotspot for methylation. For instance, MS/MS fragmentation of MMFT-8H_2_ yielded peaks corresponding to ions of MFT-1H_2_ (398.18011) and MMFT-2H_2_ (574.24640) suggesting that the methyl group is present in the second sugar moiety (Fig. 2a).

### Structure elucidation of the oligosaccharide moiety

To determine the exact structure of elongated mycofactocins we conducted large-scale fermentations (50 L) in a fermentor and harvested the resulting biomass. Cell lysis and MS-guided purification using a solid-phase extraction procedure resulted in an enriched fraction of the dominant mycofactocin. This species exhibited the same exact mass and fragmentation pattern as MMFT-2H_2_, but eluted at a slightly shifted retention time (Supplementary Fig. 3). We therefore named this species MMFT-2bH_2_.

Due to the low yields of individual MMFT derivatives, co-elution of contaminants and intrinsic purification problems associated with oligoglycosylated compounds, structural analysis by nuclear magnetic resonance (NMR) was not possible at this stage. Thus, we used enzymatic degradation to obtain structural information about the oligoglycoside chain. Gratifyingly, cellulase (*β*-1,4-glucanase) degraded the sugar chain of mycofactocin species (n>2), while amylase (α-1,4-glucanase) did not exhibit any effect (Fig. 3a). This finding strongly suggested that the oligosaccharide chain represents a *β*-1,4-glucan (Fig. 3b). Intriguingly, isomer MMFT-2bH_2_, not MMFT-2H_2_ accumulated after the enzymatic digest, suggesting that MMFT-2b(H_2_) represents a product of cellulase digestion of MMFT-n(H_2_) and shares the identical disaccharide anchor. In contrast, MMFT-2(H_2_) is a minor MMFT species found *in vivo* and might be capped by a different methyldisaccharide.

**Figure 3.**
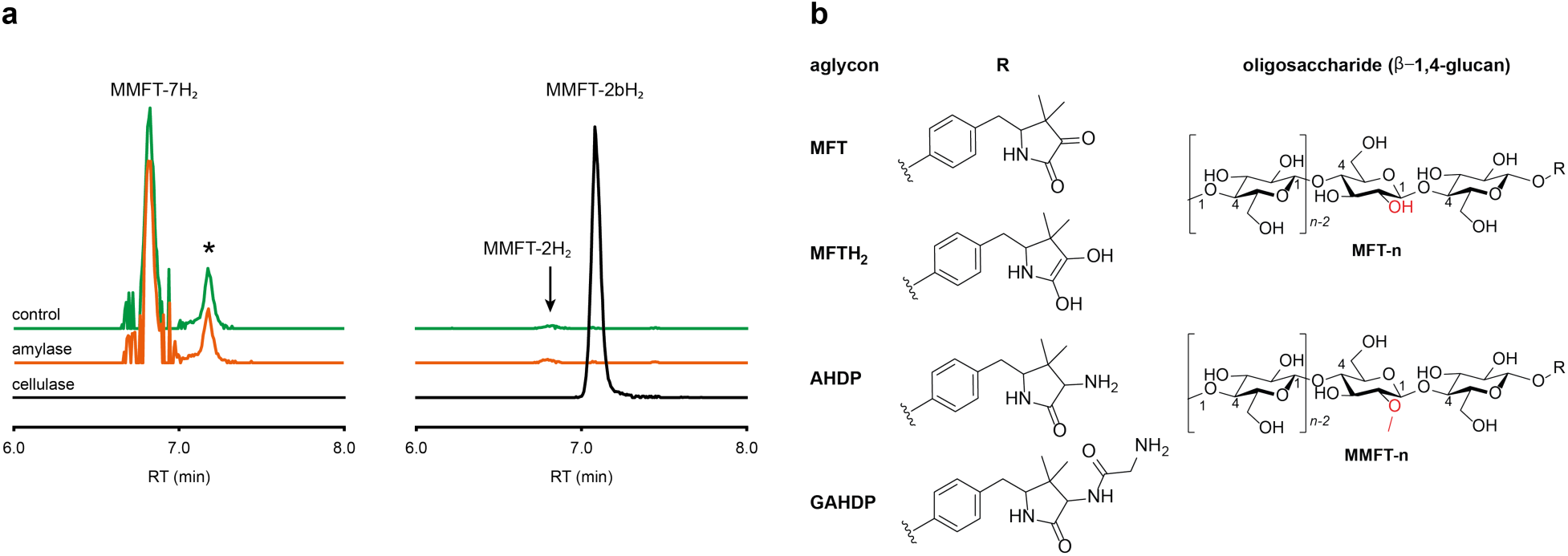
Structure of mycofactocins. **a** Enzymatic degradation of MMFT-n by cellulase. Extracted ion chromatograms (XIC, [M+H]^+^) of metabolome extract of *M. smegmatis* WT corresponding to MMFT-7H_2_ (*m/z* 1383.50626, left stack) and MMFT-2H_2_ or MMFT-2bH_2_ (*m/z* 574.24640, right stack) after treatment with cellulase, amylase, or buffer without enzyme (control) are shown. Asterisk designates a peak corresponding to the M+2 isotope of MMFT-7. Digestion by cellulase (*β*-1,4-glucanase) consumes MMFT-nH_2_ and produces MMFT-2bH_2_ suggesting that the oligosaccharide consists of *β*-1,4-linked glucose. **b** Proposed chemical structures of dominant mycofactocins and biosynthetic intermediates. Mature mycofactocins are glycosylated by sugar chains consisting of up to nine β-1,4-linked glucose units (n ≤ 9). In methylated mycofactocins (MMFT), the second hexose appears to be methylated (2-*O*-methyl-D-glucose). The aglycon is PFMT or PMFTH_2_ in mycofactocinones or mycofactocinols, respectively. The aglycon is AHDP or GADHP in biosynthetic precursors AHDP-n and GAHDP-n, respectively.

To further corroborate the structure of MMFT-nH_2_, we analyzed enriched fractions of MMFT-2bH_2_ and MMFT-nH_2_ produced in shake flasks by chemical derivatization and gas chromatography coupled with mass spectrometry (GC-MS). Monosaccharides were released by acid hydrolysis and derivatized by trimethylsilylation (TMS). By carefully investigating peaks arising from the MMFT-nH_2_ and MMFT-2bH_2_ fractions as well as different carbohydrate standards, we indeed confirmed the presence of D-glucose (Supplementary Fig. 4) and 2-*O*-methyl-D-glucose (Supplementary Fig. 5) based on the identification of two 5TMS products and two 4TMS products which exhibited identical retention times and MS spectra as 5TMS products of standard D-(+)-glucose and 2-*O*-methyl-D-glucose. This result was in agreement with the enzymatic digest and suggested that the methylated sugar present in MMFT-n(H_2_) and MMFT-2b(H_2_) is 2-*O*-methyl-D-glucose. To confirm the glycosidic linkage positions, the oligosaccharide chain was permethylated before acid hydrolysis so that only hydroxyl groups involved in glycosidic bond formation would be free for silylation^23^. This experiment (Supplementary Fig. 6) lead to the formation of glucose with 2,3,6-*O*-methyl-1,4-*O*-TMS modification (diagnostic fragment ions of *m*/*z* at 88, 133, 159), confirming the 1,4 glycosidic linkage. The same products with identical retention time and MS spectra were observed from acid hydrolysis of permethylated cellulose.

Methanolysis of the permethylated MMFT-2b and MMFT-n leading to an additional methylation on the anomeric hydroxyl group confirmed the previous assignment. Methanolysis and subsequent TMS derivatization (Supplementary Fig. 7) revealed the presence 1,2,3,6-*O-*methyl-4-*O*-TMS-glucose. The same products were observed from methanolysis of permethylated cellulose (all EI-MS spectra are shown in Supplementary Figs. 8-19). It should be noted that due to the low abundance of MMFT it was only possible to apply enriched fractions for GC-MS analysis. Thus, several peaks were observed after acid hydrolysis (Supplementary Fig. 4 and 5) that originated from an unknown methylated hexose as suggested by high similarity of the MS spectrum with 3-*O*-methyl-1,2,4,6-*O*-TMS-glucose, but eluting at a different retention time. Since this product contradicted LC-MS results and enzymatic degradation, we assumed that the major component of the MMFT-nH_2_ enriched fraction was a contamination.

In summary, we propose that the oligosaccharide moiety of MFT is a *β*-1,4-glucane (cellulose). The methylated hexose present in MMFT-n(H_2_) and MMFT-2b(H_2_) was shown to be 2-*O*-methyl-glucose. The fact that MMFT-2 and MMFT-2b (digested MMFT-n) are distinct in retention times points to some degree of structural diversity within MMFTs. Notably, cellulose was shown to be produced by *M. tuberculosis* as a constituent of biofilms after exposure to reductive stress.^24^ The production of methylated glucans, like 6-*O*-methylglucose lipopolysaccharides (MGPL), albeit with α-1,4 linkage, is well described in *Mycobacteria*^25^, 2-*O-*methylglucose appears to be less common. To the best of our knowledge, MFT and mycothiol^4^ are the only cofactors that are decorated with sugar moieties.

### Glycine-derived intermediates of MFT biosynthesis

Surprisingly, the MS/MS network (Fig. 2a) revealed two additional compounds (*m/z* 1426.54 and 1588.59)) with an unusual mass shift compared to the MFT-n(H_2_) candidates. Their mass differences and MS/MS spectra indicated that they represented hepta- and octaglycosylated species sharing a head moiety closely related to PMFT and PMFT(H_2_). The molecular masses and MS/MS spectra of the compounds could be explained by the assumption that the aglycon corresponded to glycyl-AHDP (GAHDP) and these compounds represented the oligoglycosylated forms GAHDP-7 and GAHDP-8. Since the VY core peptide of MftA is preceded by a glycine residue at its N-terminal side it appeared highly likely that the GAHDP-n species corresponded to premature cleavage products of the MftC-processed precursor peptide. To corroborate this hypothesis, we fed *M. smegmatis* cultures with a combination of fully ^13^C-labeled Gly-^13^C_2_, Val-^13^C_5_, and Tyr-^13^C_9_. Indeed, GAHDP-derived molecules underwent a mass shift of +15.05033 Da, indicating the incorporation of Gly-^13^C_2_ (+2.00671 Da) in addition to the decarboxylated Val-Tyr moiety (+13.04362 Da) (Supplementary Fig. 20). Targeted searches for GAHDP-n as well as AHDP-n (lacking the glycyl residue) and MADHP-n candidates (AHDP decorated with monomethylated oligosaccharide) revealed three series of oligoglycosylated compounds with similar retention times within each series (Fig. 4, Supplementary Table 2).

**Figure 4.**
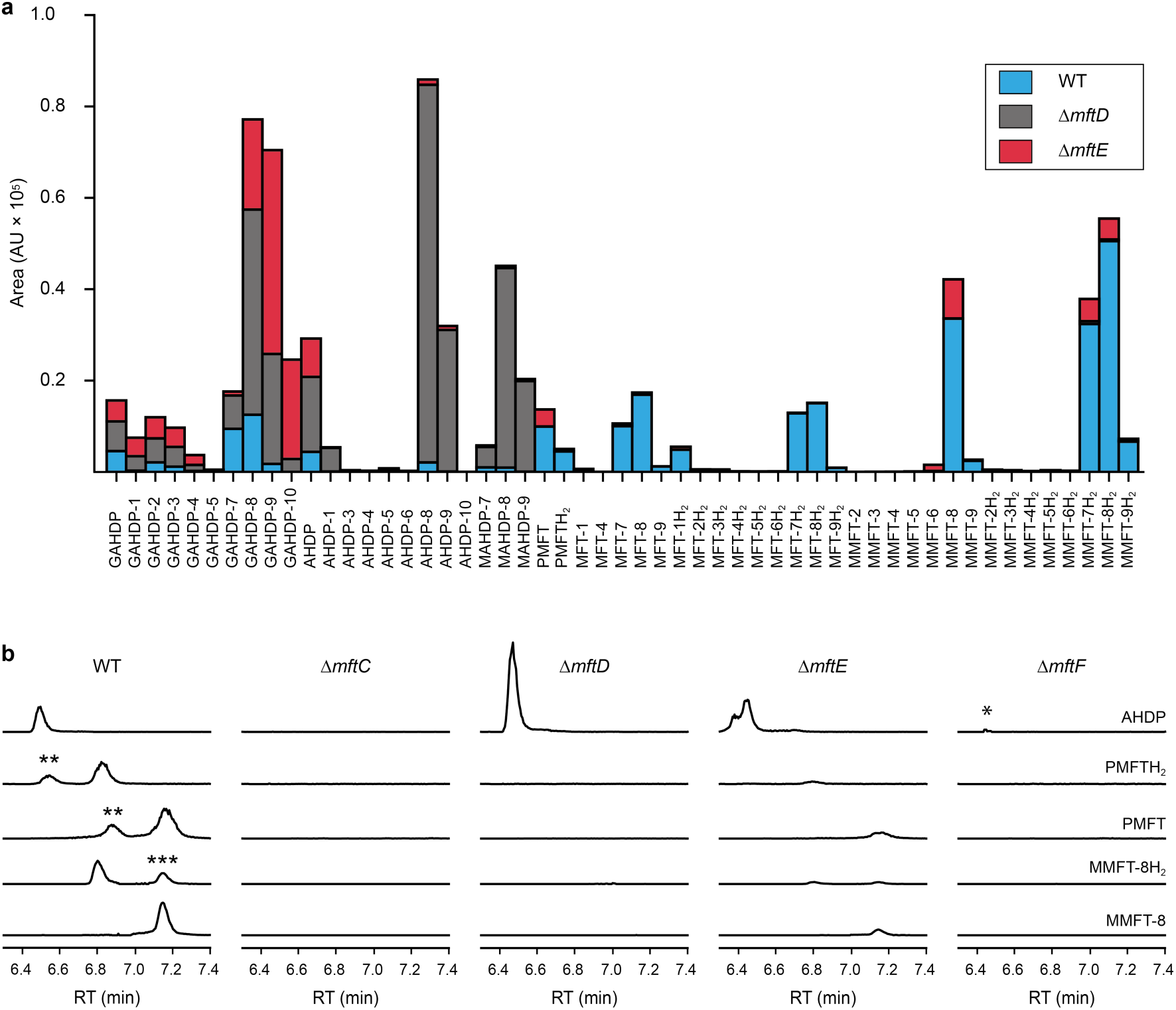
Metabolic profile of MFT congeners present in *M. smegmatis.* **a** Distribution of proposed MFT congeners as determined by HR-LCMS (Supplementary Table 2). Bars indicate area under the curve of designated species (average of three biological replicates, n = 3). Blue: WT, red: ΔmftE, gray: ΔmftD. The ΔmftE mutant produces significantly reduced amounts of MFT congeners compared to WT, but accumulates incorrectly cleaved biosynthetic intermediates (GAHDP-n series). ΔmftD is unable to produce PMFT(H_2_) and glycosylated (M)MFT-n(H_2_), thus accumulating AHDP-n congeners. **b** Extracted ion chromatograms (XIC, [M+H]^+^) of WT and mutants (ΔmftC, ΔmftD, ΔmftE, ΔmftF) corresponding to AHDP (*m/z* 235.14411), PMFT (*m/z* 234.11247), PMFTH_2_ (*m/z* 236.12812), MMFT-8 (*m/z* 1544.55072) and MMFT-8H_2_ (*m/z* 1546.56637). **marks minor isomeric forms (Supplementary Fig. 2). ***marks a peak corresponding to the M+2 isotope of MMFT-8. ΔmftC is blocked in biosynthesis of all MFT intermediates, ΔmftF abolishes most of the MFT products, but forms trace amounts of AHDP (*). ΔmftE produces most mature MFT species in lower amounts, while intermediates like AHDP are increasing. ΔmftD strongly accumulates AHDP, while MFT congeners are abolished.

### Dissection of MFT biosynthesis

In order to test if all of the MFT candidate compounds were related to MFT biosynthesis, we started to investigate mutants (ΔmftC, ΔmftD, ΔmftE, ΔmftF) of the MFT biosynthesis pathway for the production of candidate molecules. Since MFT mutants, except for ΔmftE, do not grow on ethanol as a sole source of carbon^12^, we first cultivated bacteria on the surface of cellulose filters on agar plates containing 10 g L^−1^ glucose as a carbon source. The filters were then transferred to treatment plates containing 10 g L^−1^ ethanol and incubated overnight. This strategy allowed for induction of MFT production in the wild type (WT) while ensuring comparable conditions for the mutant strains. We then compared the metabolic profile of all mutant strains (Fig. 4a, Supplementary Table 2). Indeed, none of the MFT precursors, nor any of the glycosylated candidates were detected in the ΔmftC knock-out strain (Fig. 4b). This finding, together with the fact that the genetically complemented strain ΔmftC-Comp restored production of MFT congeners (Supplementary Table 2) represented strong evidence that we indeed identified *bona-fide* MFT-derivatives.

The ΔmftE mutant was able to produce (M)MFT-nH_2_ candidates, albeit in significantly lower amounts, explaining the previously unexpected phenotypic observation that the ΔmftE mutant was able to grow on ethanol, but slower than WT^12^. Intriguingly, the pool of GAHDP-n was strongly increased in the ΔmftE strain (Fig. 4b). We thus conclude that MftE can be complemented by an unknown peptidase present in the metabolic background of mycobacteria. Theoretically, an aminopeptidase would be sufficient to degrade the N-terminus of MftA, releasing the AHDP-like core. Peptidases encoded outside the biosynthetic gene cluster have been observed in other RiPP biosyntheses as well^26^. However, the removal of the glycine residue is an apparent bottle-neck of the alternative maturation pathway in *M. smegmatis*.

In full agreement with the *in-vitro* finding that MftD consumes AHDP to form PMFTH_2_^21^, all metabolites downstream of (M)AHDP-n were abrogated in the ΔmftD strain, whereas AHDP-n and GAHDP-n accumulated (Fig. 4). The fact that GAHDP-n increased might suggest that the MftE step is impeded in the absence of MftD as well. Genetic dysregulation or cooperative effects between the two enzymes, like complex formation and substrate channeling, might account for this result.

### Glycosylation of MFT is mediated by MftF

It has been speculated that the putative glycosyltransferase MftF catalyzes a final glycosylation of PMFT to yield the mature cofactor^21^. Our results at this point showed that multiple glucose residues are attached to the aglycon *in vivo*. Furthermore, glycosylation appeared already at an early stage as mirrored by the presence of the glycosylated (G)AHDP-n series. In order to link oligoglycosylation to a given gene product, we analyzed the ΔmftF mutant for the production of glycosylated MFT congeners. Indeed, all glycosylated MFT congeners were abolished in the ΔmftF metabolome. Unexpectedly, ΔmftF mutants additionally ceased to produce the aglycons PMFT and PMFTH_2_. MftF did, however, produce trace amounts of AHDP, thus showing that at least residual MftC activity was present in the mutant (Fig. 4b). To exclude polar effects, we complemented ΔmftF by re-introduction of the *mftF* gene under control of the *mftA* promotor. The restoration of the full MFT metabolite spectrum (Supplementary Table 2) excluded polar effects and thus verified that MftF was the glycosyltransferase responsible for oligoglycosylation of MFT congeners. The appearance of glycosylated (G)AHDP species in WT together with the drastic decrease of aglycon precursors in ΔmftF can be interpreted in a scenario where MftF is a central factor of MFT biosynthesis *in vivo*. If missing, the biosynthetic machinery may fail to assemble a functional complex or may be unable to recruit the unglycosylated metabolic precursors. The finding that the *mftF* gene is a conserved constituent of MFT biosynthetic loci among different phyla supports the importance of this modification^11^. The deduced MftF protein of *M. smegmatis* (MSMEG_1426) consists of 470 amino acids (aa) and belongs to the glycosyltransferase 2 family (GT2) according to PFAM (PF00535) and CAZy searches. These enzymes are known for an inverting mechanism of oligoglycoside formation. This is well in agreement with the proposed β-configuration of the MFT oligosaccharide chain. Sequence alignment (Supplementary Fig. 21a) showed a high degree of sequence conservation among mycobacterial species and other actinomycetes (e.g., 92% similarity to MftF of *M. tuberculosis* H37Rv). Prediction of transmembrane domains revealed a single helix spanning residues 324 – 346 with the N-terminus being located outside of the membrane Supplementary Fig. 21b). The MMFT biosynthetic machinery, however, appears not to be fully encompassed within the MFT cluster since no methyltransferase was found. Future studies are warranted to identify the enzymes involved in MFT oligosaccharide methylation.

### Cofactor role of mycofactocin

After discovery of the full set of mycofactocins, we examined to which extent their production was actually dependent on the presence of ethanol in culture media. We therefore systematically compared the metabolome of *M. smegmatis* WT after ethanol treatment with glucose controls. (Supplementary Table 3). The results demonstrated that all MFT congeners and intermediates were strongly upregulated upon cultivation on ethanol (median: 34-fold upregulation). (Fig. 5a). These data perfectly support a recent report that MFT is involved in alcohol metabolism^12^.

**Figure 5.**
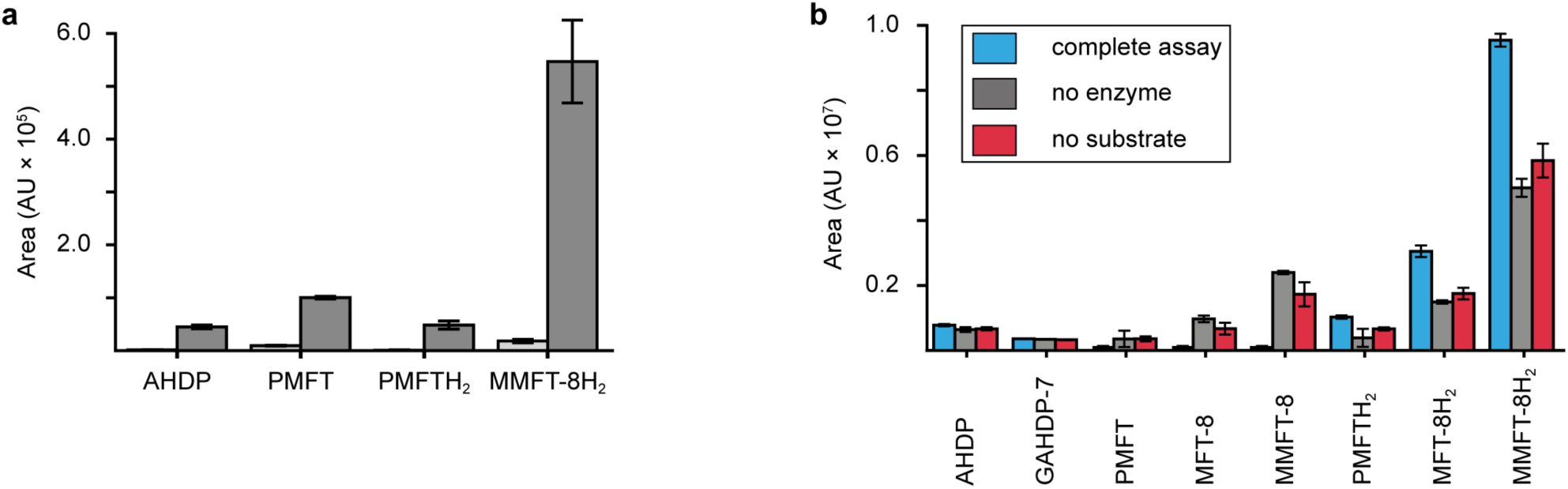
Cofactor role of mycofactocin. **a** Induction of MFT production by ethanol. Area under the curve of selected MFT precursors and congeners produced by *M. smegmatis* WT treated with ethanol (dark gray) versus glucose controls (light gray) are shown. All MFT congeners were strongly upregulated (AHDP: 33-fold, PMFT: 11-fold, PMFTH_2_: 54-fold, MMFT-8H_2_: 26-fold) on ethanol-containing media (Supplementary Table 3). Bars represent average area under the curve, error bars standard deviation of area under the curve of corresponding metabolites extracted from 3 biological replicates (n = 3). **b** Reduction of mycofactocinones to mycofactocinols by carveol dehydrogenase (activity-based metabolic profiling). The metabolome extract of *M. smegmatis* grown in ethanol-containing media was treated with recombinant L-carveol dehydrogenase LimC from *Rhodococcus erythropolis* and carveol as a substrate. Bars show average areas under the curve of selected MFT congeners resulting from enzyme treatment and controls. Error bars indicate standard deviation of 3 biological replicates (n = 3). Blue: Complete assay with enzyme and L-carveol as substrate. Red: Control without substrate, Gray: Control without enzyme. Mycofactocinones (PMFT, MFT-8, MMFT-8) were completely depleted in assay conditions, while levels of mycofactocinols (PMFTH_2_, MFT-8H_2,_ MMFT-8H_2_) increased ca. 2-fold compared to no-enzyme controls. Slightly higher levels of mycofactocinols in controls lacking substrate compared to controls lacking enzymes can be explained by reduction by enzyme-bound NADH. Levels of redox-inactive precursors (AHDP and GAHDP) remained unaltered under all assay conditions.

Finally, we sought to confirm that the MFT congeners identified in this study are actually coenzymes of MFT-dependent enzymes. To assess this question, we turned to activity-based metabolic profiling^27^. We incubated the extracted metabolome of *M. smegmatis* with the recombinant carveol dehydrogenase LimC from *Rhodococcus erythropolis* (Supplementary Fig. 22), a nicotinoprotein with a non-exchangable NADH cofactor^28^. This enzyme was proposed to require MFT as an external electron acceptor^11^. A recent study showed that carveol dehydrogenase from *M. smegmatis* was able to reduce PMFT to PMFTH_2_ using carveol and internally bound NADH as an electron donor^21^. Likewise, we observed full reduction of all mycofactocinones to mycofactocinols (Fig. 5b, Supplementary Table 4) by LimC when combined with carveol as a substrate. Controls lacking enzyme or substrate showed weak and no turnover, respectively. The low turnover by LimC alone can be explained by internally bound NADH as reported before^21^. Both the aglycon PMFT as well as the oligoglycosylated MFT-n and MMFT-n species were completely turned over, while redox-inactive AHDP congeners remained unaffected. These data further validate the notion that all MFT candidates presented here are *bona-fide* mycofactocins with full cofactor function.

## Conclusion

The proposed redox cofactor mycofactocin has attracted considerable interest since it was postulated by bioinformatics. Despite recent progress made by *in-vitro* studies, evidence for mycofactocin congeners in living microorganisms has been missing so far. Our metabolomics approach combined with stable isotope labeling, induction by ethanol, as well as genetic dissection of the biosynthetic pathway turned out to be a powerful approach to identify MFT congeners in mycobacteria. We discovered that MFT is decorated with oligosaccharides consisting of up to nine β-1,4-linked glucose units. Analyses of ΔmftF mutants and complement strains revealed that MftF is the glycosyltransferase responsible for the oligoglycosylation observed. Mycofactocins can be isolated in oxidized (mycofactocinones) and reduced forms (mycofactocinols) and are co-substrates of enzymatic reduction by carveol dehydrogenase. These data provide strong evidence that mycofactocins are indeed redox cofactors as proposed earlier^11, 12, 21^. We therefore conclude that we have finally discovered the family of compounds that was tentatively called "mycofactocin" and thus close an important gap of knowledge in the field. Our results will guide further studies into the occurrence, physiological role, and biochemistry of mycofactocins in microorganisms. Finally, we are confident that this work will inspire future efforts to exploit mycofactocin as a disease marker or as a potential drug target for the treatment of tuberculosis and other mycobacterial infections.

## Methods

### Microbial strains and general cultivation conditions

The *M. smegmatis* MC^2^ 155 WT and mutant strains ΔmftC, ΔmftD, ΔmftE and ΔmftF were maintained in lysogeny broth (LB), supplemented with 0.05% Tween80 (LB-Tween) at 37 °C and 180 rpm. Complement mutants *M. smegmatis mftC*-Comp and *mftF*-Comp were maintained in media containing kanamycin (20 µg mL^−1^). *Escherichia coli* TOP10 was grown in LB supplemented with kanamycin (30 µg mL^−1^) at 37 °C for the generation and maintenance of pMCpAINT-derived plasmids. Metabolomics studies were carried in adapted HdB medium^29^ containing 3 g L^−1^ Na_2_HPO_4_. Liquid cultures were supplemented with 0.5 g L^−1^ tyloxapol, 10 g L^−1^ glucose (w/v) or 10 g L^−1^ ethanol (w/v) at 37 °C and 180 rpm.

### *Generation of M. smegmatis* mutant and complement strains

Scarless mutants of biosynthetic genes of *M. smegmatis* (ΔmftC, ΔmftD ΔmftE, ΔmftF) as well as the *mftC* complement strain (ΔmftC-Comp) were obtained from the Kaufmann laboratory^12^. For the genetic complementation of *M. smegmatis ΔmftF*, the promotor and the ribosome binding site of *mftA* were combined with the *mftF* coding sequence (obtained as a synthetic DNA construct) and cloned as an insert of the integrative plasmid pMCpAINT^30^ yielding plasmid pPG20 (Supplementary Fig. 23). After transformation of electrocompetent cells of *M. smegmatis* by electroporation, positive clones (ΔmftF-Comp) were selected on kanamycin (20 µg mL^−1^). Complement mutants were confirmed phenotypically by the (restored) ability to grow in HdB supplemented with 10 g L^−1^ ethanol as a sole source of carbon as well as genetically by positive PCR amplification of the *mftF* gene using the following primers: INT_mftF_F: 5’-ACTTCTCCGGTATGCACTGC-3’ and INT_mftF_R1: 5’-ACAGATCGCCGAACACAACT-3’).

### Isotopic labeling of M. smegmatis

A saturated pre-culture of *M. smegmatis* MC^2^ 155 WT in LB broth was used to inoculate 25 mL of HdB medium supplemented with 0.5 g L^−1^ tyloxapol and 10 g L^−1^ ethanol to an initial optical density at 600 nm (OD_600_) of 0.1. Cultivations contained 1 mM L-tyrosine-^13^C_9_ (99% atom purity, Cortecnet) and 1 mM L-valine-^13^C_5_ (99% atom purity, Merck) and were conducted in four replicates at 37 °C and 180 rpm for 24 h. Cultures (10 mL of a 10-fold dilution) were poured onto sterile regenerated cellulose filters (0.2 µm, Sartorius) previously conditioned with water. The biomass was repeatedly washed with sterile water and transferred to HdB agar plates with 10 g L^−1^ ethanol and with either light or heavy L-valine and L-tyrosine supplementation, as appropriate. Filters inoculated with sterile media were used as control. Filters were incubated at 37 °C for 48 h. Directly after the incubation period, the filters were extracted and subjected to LC-MS measurements as described below. Data analysis was performed with the Stable Isotope Labeling workflow of Compound Discoverer 3.0 (Thermo Scientific) allowing for a maximum exchange of 16 ^13^C atoms. Independent analyses were performed for the lower and higher scan ranges. A minimal peak intensity cut-off of 10^4^ was defined for compound detection. Compounds with the same mass (± 5 ppm) and eluting within 0.2 min from each other were grouped. Molecules were considered candidates potentially comprising the decarboxylated Val-Tyr core peptide (*i.e.* AHDP moiety) if a relative ^13^C-exchange rate higher than 50% was observed. Low abundance compounds (area <1000) and candidates containing a high proportion of contaminating masses were disregarded.

### Comparative metabolomics studies

Cultures of *M. smegmatis* MC^2^ 155 WT as well as ΔmftC, ΔmftD, ΔmftE, ΔmftF, ΔmftC**-**Comp and ΔmftF-Comp growing in LB supplemented with 0.05% of Tween80 were used to inoculate sterile regenerated cellulose filters as described before, standardizing all the cultures to the same concentration. The filters were incubated in HdB supplemented with 10 g L^−1^ glucose at 37 °C for 18h. Afterwards, the filters were transferred to a new HdB plate supplemented either with 10 g L^−1^ of glucose or 10 g L^−1^ ethanol and incubated at 37°C for 18h. This study was carried out in triplicates and filters incubated with media were used as control (blank). Directly after the incubation period, the filters were further extracted and subjected to LC-MS measurements as described below. Targeted studies were performed using the Expected Compounds node in Compound Discoverer 3 with an Expected Compounds table including AHDP, PMFT, PMTH_2_, MFT-1 and MFT-1H_2_ allowing for multiple glycosylation and methylation events. Compounds from different runs with the same mass (< 5 ppm deviation) and eluting within 0.2 min from each other were grouped. Median of areas under the curve of three replicates was used to compute ratios between groups.

### Metabolite extraction and LC-MS measurements

Filters were recovered with sterile tweezers and placed on 20 mL chilled extraction mixture (acetonitrile:methanol:water 60:20:20 with 0.1% formic acid). Bottles were placed one hour at −80 °C, sonicated for 5 min in an ultrasonic bath at room temperature and frozen. The lysis procedure was repeated three times. After the last ultrasound treatment, extract was transferred to round flasks, frozen and lyophilized. The dry extract was resuspended in 950 µL ddH_2_O, centrifuged twice and the final extract was saved in HPLC vials at −20 °C until measurement. LC-MS measurements were performed on a Dionex Ultimate 3000 system combined with a Q Exactive Plus mass spectrometer (Thermo Scientific) equipped with a heated electrospray ion source (HESI). Metabolite separation was carried out using a Phenomenex Kinetex XB-C18 column (150 × 2.1 mm, 2.6 µm, 100 Å) preceded by a Phenomenex SecurityGuard ULTRA guard cartridge (2 × 2.1 mm). Mobile phases consisted of 0.1% formic acid in either water (A) or acetonitrile (B). 10 µl of the sample were separated chromatographically at 40 °C and a constant flow rate of 300 µL min^−1^ as follows: 0-2 min, 2% B; 2-15 min 2-99% B; 15-18 min 99% B. Metabolite separation was followed by both full scan (MS^1^) and data-dependent MS/MS (MS^1^ and Top10 MS/MS) analyses in positive ionization mode at two scan ranges: *m/z* 200 to 600 and *m/z* 580 to 2000. Spray quality was adjusted based on the ratio of water:acetonitrile. For the first 10 min of elution sheath gas flow rate was set to 35 and auxiliary gas flow rate to 7. For the following 12 min those values were set to 24 and 2, respectively. Capillary temperature was 320 °C, probe heater temperature was 230 °C, spray voltage was 4 kV, and S-lens RF level was 50 at all times. MS^1^ had the resolving power set to 70,000 FWHM at *m/z* 200, injection time to 100 ms, and AGC to 3E6. The ten most intense ions were selected for MS/MS with a scan rate of 12 Hz with a dynamic exclusion of 10 s. Resolving power was 17,500 FWHM at *m/z* 200, AGC target was 1E5, and injection time was 50 ms. Isolation window was set to *m/z* 1, while routine analysis was performed at 40 NCE (normalized collision energy). Targeted MS/MS spectra acquisition for the MFT congeners was performed at NCE values of 20, 30, and 40.

### MS/MS networking

*M. smegmatis* MC2 155 metabolome extracts were prepared as described before. Data-dependent (TOP 10) MS/MS data was acquired and converted to the mzXML format with the MSConvert tool^31^. Files were uploaded to the Global Natural Products Social Molecular Networking (GNPS) server^22^ and analyzed with the Molecular Networking pipeline. Parameters were set as default except for the following: minimum cosine score (0.6) and minimum fragment ions (4). Results were visualized in Cytoscape 3.7^32^.

### Bioinformatics analysis of the MftF primary structure

The primary protein structure of several MftF homologues was downloaded from the NCBI database. Sequences were aligned using the MUSCLE algorithm^33^ implemented in Geneious Prime (2019.1.1). Prediction of transmembrane domains was performed using the TMHMM 2.0 webserver^34^. Classification of MftF was performed by the carbohydrate-active enzymes database (CAZy)^35^.

### Heterologous production and purification of LimC

The *limC* gene encoding the carveol dehydrogenase LimC^28^ from *Rhodococcus erythropolis* DCL14 was obtained as a codon optimized synthetic construct inserted in vector pET28, in-frame with the N-terminal hexahistine tag (plasmid pLAPO4, see Supplementary Fig. 23a). A single colony of *E. coli* BL21(DE3) freshly transformed with plasmid pLAPO4 was inoculated in 5 mL LB with kanamycin (50 µg mL^−1^) and cultured overnight at 37 °C and 210 rpm. A main culture (100 mL) was inoculated in LB media amended with the same antibiotic and cultured until an OD_600_ of 0.5 – 0.6. At this point expression of *limC* was induced with IPTG (0.5 mM) and the temperature decreased to 16 °C, then further cultured for 18 – 20 h. Cells were harvested from the medium by centrifugation at 4000 × g and 4 °C for 30 min, then resuspended and washed in cold sodium chloride 0.85% and harvested again for 15 min, then frozen at −20 °C for future use or processed as follows. The cell pellet was resuspended in lysis buffer (50 mM NaH_2_PO_4_, 300 mM NaCl, pH 8.2) containing 20 mM imidazole, disrupted with a Sonopuls ultrasonic sonifier (Bandelin), and centrifuged at 17000 × g, 4 °C for 30 min. The lysate was applied to 1.25 mL HisPur NiNTA resin (Thermo Fisher Scientific), pre-equilibrated with 10 column volumes of lysis buffer at the same imidazole concentration and allowed to flow at room temperature by gravity. The resin was washed with 10 CV of wash buffer (lysis buffer containing 30, 50, and 70 mM imidazole). The target protein was then eluted with 2.5 mL of lysis buffer containing 300 mM imidazole, then immediately rebuffered in 3.5 mL assay buffer (sodium citrate buffer 50 mM, pH 6) in a PD-10 gel filtration column (GE Healthcare) and concentrated to a final volume of 500 µL in a 3 kDa MWCO Vivaspin centrifugal filtration unit (Sartorius). The protein was detected by SDS-PAGE (Supplementary Fig. 22), protein concentration was determined with the Roti Nanoquant (Carl Roth) reagent following manufacturer instructions in relation to known concentrations of BSA in assay buffer. Positive enzymatic activity was confirmed by incubating with L-carveol (1 mM) in the presence of 0.1 mM DCPIP. Discoloration in the presence of carveol indicated positive activity.

### Activity-based metabolic profiling

Metabolome extracts of *M. smegmatis* MC^2^ 155 were prepared as described above and used for activity-based metabolic profiling. Metabolome extract (10 µL of a 10-fold dilution) was mixed with 0.1 µg of purified LimC in assay buffer (sodium citrate buffer 50 mM, pH 6) containing L-carveol (1 mM) in a 20 µL reaction volume. Negative controls were performed without substrate or without enzyme. The reaction was quenched after 1 h of incubation at room temperature by adding 1 volume of LC-MS grade acetonitrile and centrifugation for 10 min at 4°C. 10 µL were injected for LC-MS measurement as described below. Mycofactocinone and mycofactocinol species were identified using Compound Discoverer 3.1 based on *m/z* and retention time.

### Glucanase digestion of mycofactocins

Metabolome extracts of *M. smegmatis* MC^2^ 155 were prepared as described above and mixed with 0.2 mU of commercial cellulase (1,4-β-glucanase) from *Trichoderma reesii* ATCC 26921 (Merck) in 50 mM citrate-phosphate buffer at pH 5 (6.5 mM citric acid, 43.6 mM sodium phosphate) or 0.2 mU α-amylase (1,4-α-glucanase) from *Bacillus licheniformis* (Merck) in citrate-phosphate buffer at pH 7 (24.3 mM citric acid, 25.7 mM sodium phosphate). Enzyme solutions were prepared at a concentration of 0.2 mU µL^−1^. 30 µL reactions were set up with 1 µL enzyme, 14 µL of metabolome extract and 15 µL of concentrated buffer (2 ×). Buffer replaced the enzyme solution at pH 5 or 7 in control measurements. The reaction was incubated for 1 hour at the optimum temperature for each enzyme, then quenched by the addition of 1 volume acetonitrile (LC-MS grade) and centrifuged at 12000 × g for 10 min. 5 µL of the supernatant were injected for LC-MS measurements.

### Enrichment of the MMFT-2b(H2)

The first pre-culture of *M. smegmatis* WT was grown for 48 h in serum bottles under the following conditions: shaking frequency of 180 rpm, shaking diameter of 25 mm and a flask volume of 50 ml with 20 ml filling volume. The second pre-culture step was conducted in Erlenmeyer shake flaks for 48 h under the following cultivation conditions: shaking frequency of 180 rpm, shaking diameter of 25 mm and a flask volume of 2 L with a filling volume of 400 mL. The transferred volume of the second pre-culture was 1 L to inoculate a 75 L stirred tank reactor filled with 50 L LB medium supplemented with 20 g L^−1^ ethanol and 20 g L^−1^ glucose for 23.25 h of cultivation. To ensure aerobic conditions, the dissolved oxygen tension was controlled by the stirring rate and therefore always higher than 20%. The gas flow rate was constant at 0.5 vvm as well as the headspace overpressure of 0.2 bar. For biomass separation, the overall fermentation broth was filtered and the received cells were extracted twice with MeOH. The MeOH extract was separated from the cell debris by filtration and concentrated under reduced pressure. LC-MS analysis of crude MeOH extract revealed the presence of MMFT-2bH_2_ with the major detectable mycofactocin species, with the observed molecular ion at *m*/*z* 574.24945 ([M+H]^+^), and MMFT-2b in trace amounts, with the observed molecular ion at *m*/*z* 572.23431 ([M+H]^+^). The concentrated MeOH extract was first suspended into 10% MeOH (50 mL) by ultrasonication, then fractionated by SPE-C18 cartridge (10 g) with the step elution by 10% MeOH, 20% MeOH, 30% MeOH, 40% MeOH, 50% MeOH, 60% MeOH, 80% MeOH, and pure MeOH, and concentrated under reduced pressure. LC-MS analysis of individual fractions indicated that the MMFT-2bH_2_ was accumulated in 30% MeOH fraction, with a small amount in 40% ‒ 60% MeOH. The 30% MeOH fraction, as well as 40% ‒ 60% MeOH fractions were submitted to size exclusion by Sephadex LH20 eluted with 50% MeOH, and fractionated into six fractions. Following the LC-MS analysis of each fraction, the MMFT-2H_2_ molecular ion was located in the third fraction (Fr.3-3). In the end the Fr.3-3 was submitted to semiprepative HPLC coupled with Phenomenex Luna C8 column (100 Å 250 × 10 mm) and separated under the gradient of 0 ‒ 5 min, 10% MeOH; 5 ‒ 20 min, 10% ‒ 60% MeOH; 20 ‒ 25 min, 60% MeOH; 25 ‒ 35 min, 60% ‒ 100% MeOH; 35 ‒ 40 min, 100% MeOH, with the flow rate of 2.0 mL min^−1^. LC-MS indicated that MMFT-2bH_2_ was enriched in the fraction with *t*_R_ = 22.6 min, and the mixture of MMFT-2bH_2_ and MMFT-2b was enriched in next fraction with *t*_R_ = 23.0 min.

### Enrichment of MMFT-n

For large-scale cultivation, pre-cultures of *M. smegmatis* WT in LB-Tween were used (1:200) for inoculation of batches of 400 mL in 1 L flasks of LB-Tween supplemented with 10 g L^−**1**^ ethanol. In total, 24 L culture broth was inoculated, centrifuged, and bacterial pellet was collected and frozen at ‒80 °C before extraction in batches. 200 mL of MeOH was added into a glass flask containing thawed bacteria and stirred at 4 °C overnight. The yellowish MeOH extract was filtrated and concentrated under reduced pressure, suspended into 10% MeOH by ultrasonication and finally loaded on a SPE C18 cartridge (5 g). The fractionation was performed by step elution with 10% MeOH, 20%, 30%, 40%, 50%, 60%, 80%, 100% MeOH. LC-MS analysis of individual fractions indicated that the MMFT-n was accumulated in 30% MeOH and 40% MeOH fractions. These two fractions were submitted to size exclusion by Sephadex LH20 eluted with 50% MeOH, and fractionated into six fractions, respectively. Based on the LC-MS analysis of each fractions, the MMFT-n(H_2_) enriched fraction was further purified by semipreparative HPLC. MMFT-n(H_2_) containing fractions (*t*_R_ = 22.48 min) were pooled and submitted to sugar composition analysis mediated by chemical degradation, modification and finally by GC-MS analysis.

### Characterization of sugar composition

See Supplementary Information, Section 1.

### Gas chromatography - mass spectrometry (GC-MS)

GC-MS analysis was conducted using an Agilent 6890 Series gas chromatograph coupled to an Agilent 5973 quadrupole mass selective detector (interface temp, 270 °C; quadrupole temp, 150 °C; source temp, 230 °C; electron energy, 70 eV). Compounds were separated using a ZB5 column (Phenomenex, Aschaffenburg, Germany, 30 m × 0.25 mm × 0.25 µm) and He (1.5 ml min^−1^) as carrier gas. The sample (1 µL) was injected with a split of 50:1 at an initial oven temperature of 100 °C. The temperature was held for 2 min and then increased to 250 °C with a gradient of 7 °C min^−1^, and then further increased to 330 °C with a gradient of 100 °C min^−1^ and a hold of 3 min. Compounds were identified by comparison of retention times and mass spectra to those of authentic standards or by reference spectra in the Wiley and National Institute of Standards and Technology (NIST) libraries.

## Supporting information

Supplementary Table 1 - M_smegmatis 13C labeled compounds per file

Supplementary Table 2 - MFT candidates - mutants and complement

Supplementary Table 3 - MFT candidates EtOH versus glucose

Supplementary Table 4 - Activity-based metabolic profiling

## Acknowledgements

We would like to thank the Carl-Zeiss Foundation and the Leibniz Association as well as the European Regional Development Fund for financial support. CB was kindly supported by the CRC ChemBioSys 1127 (Deutsche Forschungsgemeinschaft). We thank Stefan Kaufmann and Gopinath Krishnamoorty for kindly providing *M. smegmatis* mutants.

## Conflict of interest

The authors declare that they have no conflicts of interest with the contents of this article.

## Author contributions

LP, PG, performed metabolomics experiments. LP, DB and GL analyzed metabolomics data. PG performed genetic studies. LP performed enzyme assays. LR planned and supervised the scale up in the pilot-scale reactor. HG and TK performed and analyzed GC-MS experiments. CB designed research, analyzed data and edited the manuscript. GL designed the study, analyzed data and wrote the manuscript together with DB.

## Data availability

Raw LC-MS data are available on request from the corresponding author.

## Supplementary Tables

**Supplementary Table 1: ^13^C-labeled compounds**: Table of labeled compounds (per file) computationally derived from raw LC-MS runs of bacterial extracts by Compound Discoverer 3.0. Labeled cultures were grown on media supplemented with on L-Val-^13^C_5_ and L-Tyr-^13^C_9_, unlabeled cultures grown on natural L-Val and L-Tyr. Sheet 1: candidate compounds after filtering. Sheet 2: all compounds from LC-MS with a scan range of 200-600 *m/z*. Sheet 3: all compounds from LC-MS with a scan range of 580-2000 *m/z*. Low-intensity compounds (area < 1000) are not shown. Relative exchange rates for 1-16 carbons are shown in columns J – Z.

**Supplementary Table 2**: **Comparative metabolomics comparing *M. smegmatis* WT, mutant and complement strains.** Table lists all expected MFT candidate compounds detected by LC-MS in extracts of intracellular contents. Compounds were computationally derived from raw data using Compound Discoverer 3.0 by a targeted search with indicated parent compounds. LC-MS runs with a scan range of 200-600 *m/z* and a scan range of 580-2000 *m/z* are combined in one table. Compounds shown comprise glycosylated MFT-derivatives: (P)MFT-n (mycofactocinones) (P)MFT-nH_2_ (mycofactocinols) as well as methylmycofactocinones (MMFT-n) and methylmycofactocinols (MMFT-nH_2_) with n indicating the number of sugar residues. Furthermore, proposed MFT biosynthetic intermediates, namely, AHDP-n, glycyl-AHDP-n (GAHDP-n) and methyl-AHDP-n (MAHDP-n) with n indicating the number of sugar residues are shown. Sheet 1: Compounds isolated from mutant strains (ΔmftC, ΔmftD, ΔmftE, ΔmftF) compared to WT. Sheet 2: Compounds isolated from complement strains (ΔmftC-Comp*, ΔmftF-Comp*) compared to WT.

**Supplementary Table 3: Comparative metabolomics of *M. smegmatis* WT treated with glucose or ethanol**. Table lists dominant MFT candidate compounds in extracts of intracellular contents detected by LC-MS for glucose and ethanol treatment. Compounds were computationally derived from raw data using Compound Discoverer 3.0 by a targeted search with indicated compounds. Ratios of group areas show strong increase of MFT congeners after ethanol treatment. Compounds with areas >10000 AU are shown. LC-MS runs with a scan range of 200-600 *m/z* and a scan range of 580-2000 *m/z* are combined in one table.

**Supplementary Table 4: Activity-based metabolic profiling**. Dominant MFT candidate compounds in metabolome extracts detected by LC-MS after treatment with carveol dehydrogenase LimC from *Rhodococcus erythropolis*. Compounds were computationally derived from raw data using Compound Discoverer 3.0 by a targeted search with indicated parent compounds. Enzyme treatment with LimC and carveol as substrate show cofactor function of mycofactocins: Mycofactocinons were completely depleted (ratio assay/control = 0) while the content of mycofactocinols doubled (ratio assay/control ≈ 2). Redox-inactive precursors remained unaltered (ratio assay/control ≈1). Compounds with areas >10000 AU are shown only.

