## Supplementary material for "Biosynthesis of the redox cofactor mycofactocin comprises oligoglycosylation by MftF in *Mycolicibacterium smegmatis*": Supplemetary Information

**Table of Contents**

**1 Supplementary Methods** **2**

**2 Supplementary Figures** **3**

**3 Plasmid Sequences** **26**

### 1 Supplementary Methods - Characterization of sugar composition

**Permethylation**

The enriched MMFT-n($H_2$) and MMFT-2b($H_2$) (50 μg) were transferred into a screw cap glass vial (4 mL), and dried completely in an evaporator (GeneVac). An amount of 0.5 mg of authentic standard cellulose was weighed into a 4 mL screw cap glass vial individually. To each of these samples, 0.5 mL of dried dimethyl sulfoxide (DMSO) was added together with 100 μL of iodomethane. To this solution 60 mg of finely ground NaOH was added in excess. (The pellets of NaOH were ground in a hot and dried mortor and pestle (120 °C oven for 30 min). The reaction was performed on a shaker at 50 °C for 30 min, and quenched by the addition of ice-cold water to prevent a high temperature and degradation. A liquid-liquid extraction using dichloromethane (DCM) and subsequenct washes with ice-cold water were performed three times to remove NaOH and DMSO. The upper aqueous layer was discarded, and the remaining bottom organic layer containing permethylated products was collected and dried to completion.

**Acid hydrolysis**

50 μg of the enriched MMFT-n($H_2$) and MMFT-2b($H_2$), or permethylated products were transferred into a 4 mL screw cap glass vial and dried completely by GeneVac. The acid hydrolysis was performed by adding 1 mL of HCl (3 M, aq.) and kept at 95 °C for 5 hours. Afterwards, the whole mixture was dried thoroughly by GeneVac.

**Methanolysis**

50 μg of the enriched MMFT-n($H_2$), MMFT-2b($H_2$), or the permethylated products were transferred into 4 mL screw cap glass vial and dried completely by GeneVac. The methanolysis were performed by adding 1 mL of HCl in MeOH (1.25 M) and kept at 95 °C for 5 hours. Afterwards, the whole mixture was dried thoroughly by GeneVac.

**Silylation and GC-MS**

The released monosaccharide residues were derivatized with MSTFA (MSTFA: *N*-methyl-*N*-(trimethylsilyl)trifluoroacetamide) by adding 25 μL dried pyridine and 25 μL MSTFA and kept at 60 °C for 30 min. Silylated samples were subjected to analysis by GC-MS coupled with a ZB5 column as described in the Methods section of the main text (gas chromatography - mass spectrometry). The authentic standard monosaccharides (D-(+)-glucose, methyl-α-D-glucose, methyl-β-D-glucose, 2-OMe-D-(+)-glucose, 3-OMe-D-glucose, 4-OMe-D-glucose, and 6-OMe-D-glucose) were purchased from Sigma-Aldrich or Carbosynth, derivatized and analyzed under the same procedure.

### 2 Supplementary Figures

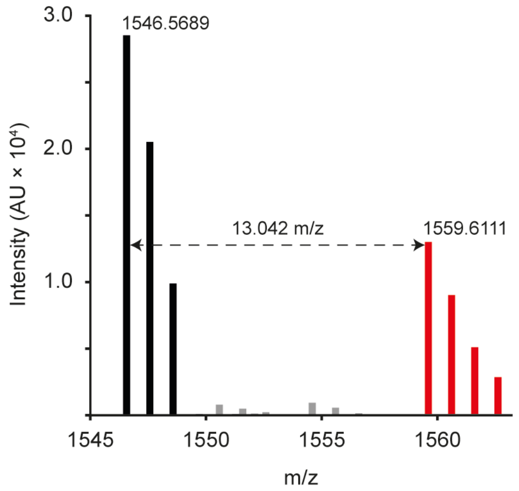


**Supplementary Figure 1. Stable isotope labeling of MFT congeners**.

After feeding of L-Val-$^{13}C_5$ and L-Tyr-$^{13}C_9$ the mass spectrum of MMFT-8$H_2$ (*m/z* 1546.56809, $[M+H]^+$) demonstrated incorporation of 13 $^{13}C$ labels (expected mass shift: 13.04362 Da). Black: Natural isotope distribution of metabolites. Red: Isotope pattern of labeled metabolite. Figure represents an overlay of two individually recorded mass spectra.

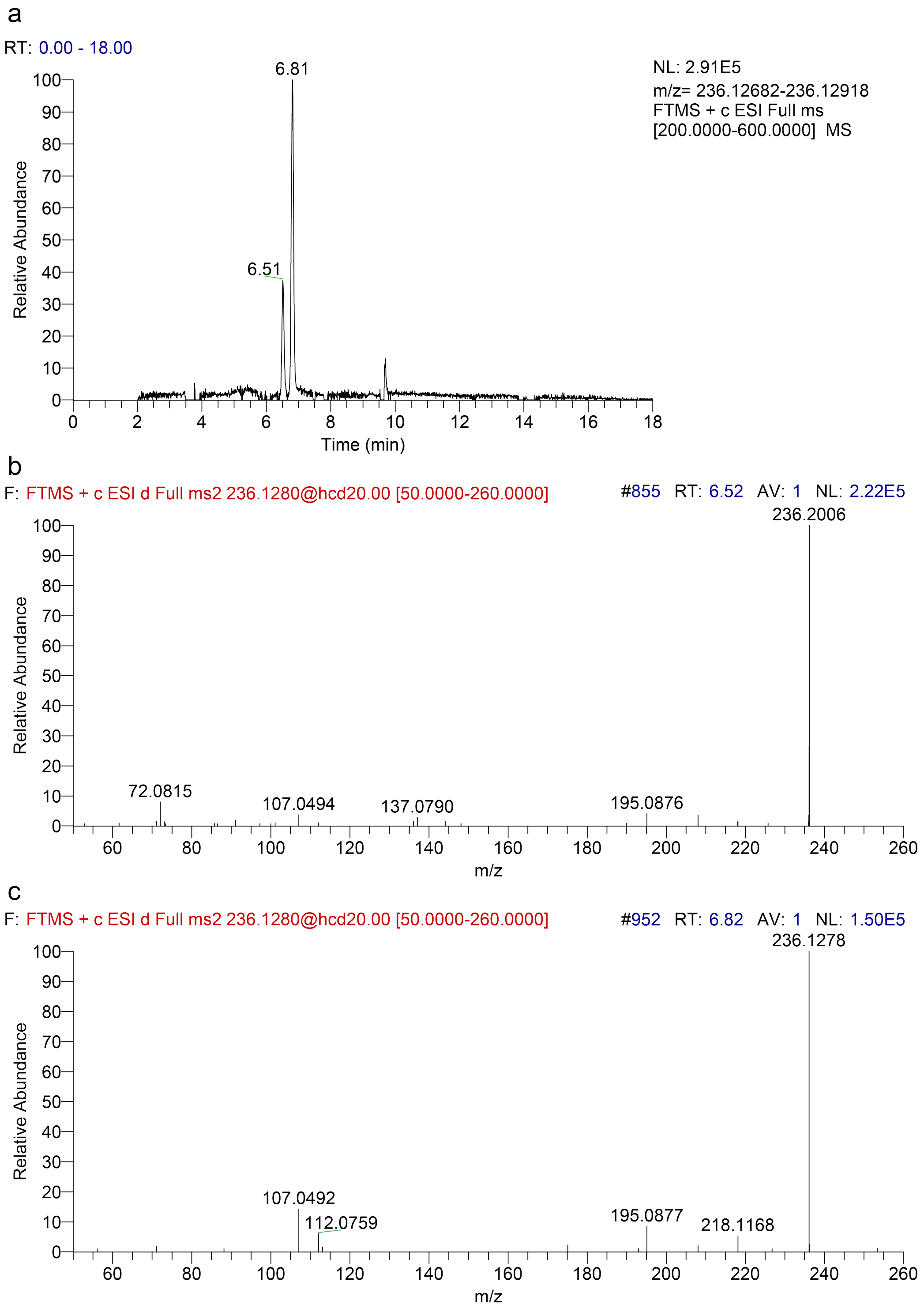


**Supplementary Figure 2. Extracted ion chromatogram of two isomeric forms of $PMFTH_2$.**
**a** Extracted ion chromatogram corresponding to $PMFTH_2$ (*m/z* 236.12812 $[M+H]^+$), showing two isomeric forms eluting at 6.51 (minor form) and 6.81 min (major form). **b** MS/MS spectrum of the minor isomer eluting at 6.52 min. **c** MS/MS spectrum of the dominant isomer eluting at ca 6.82 min. Near identical mass and MS/MS fragmentation and similar retention time suggest that the two isomers represent tautomers of each other.

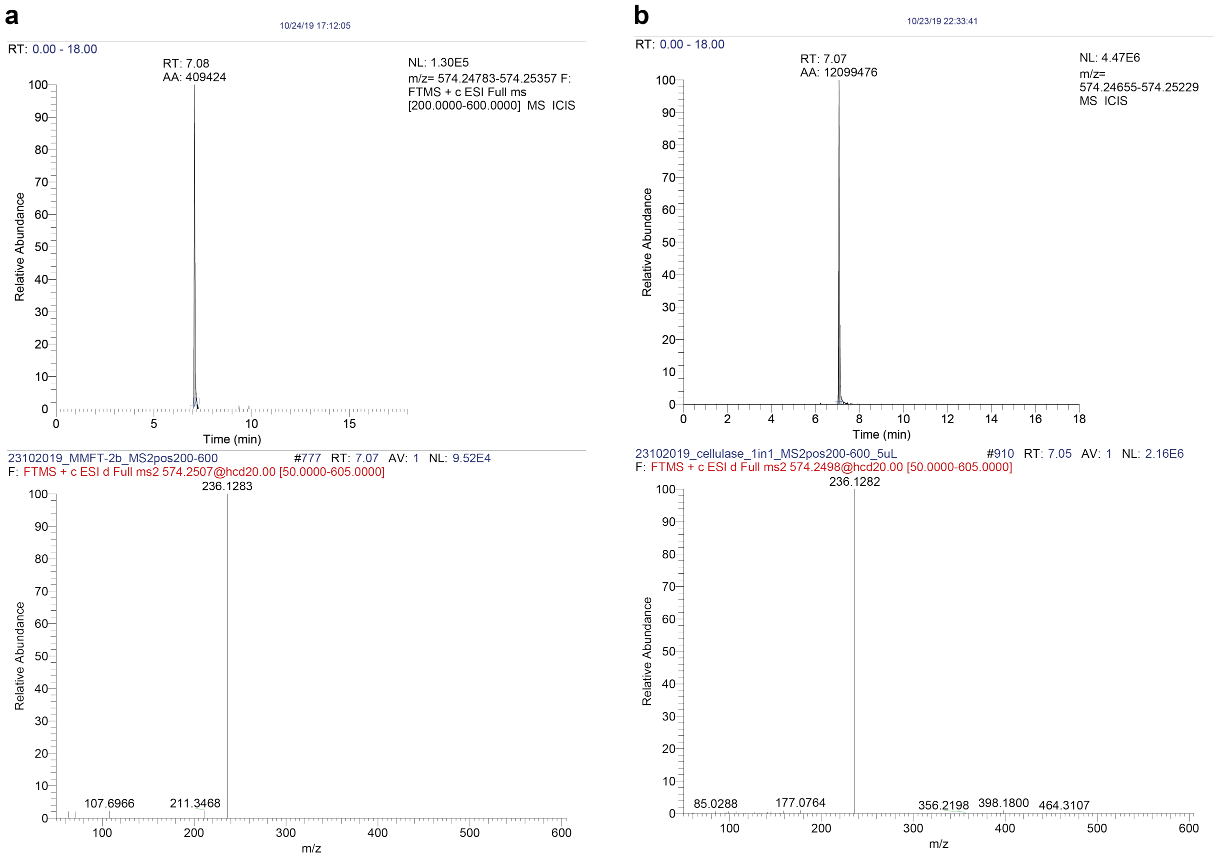


**Supplementary Figure 3. Extracted ion chromatogram and mass spectrum of MMFT-2bH$_2$.**
**a** MMFT-2bH$_2$ enriched from a large-scale cultivation (50 L) performed in a fermentor. The extracted ion chromatogram shows MMFT-2bH2 ($[M+H]^+$ *m/z* 574.24640). The corresponding MS/MS spectrum is shown below. **b** MMFT-2bH$_2$ generated by cellulase treatment of mycofactocin extracts. The extracted ion chromatogram showns MMFT-2bH2 ($[M+H]^+$ *m/z* 574.24640). The corresponding MS/MS spectrum is shown below. Both compounds are identical according to LC-MS/MS.

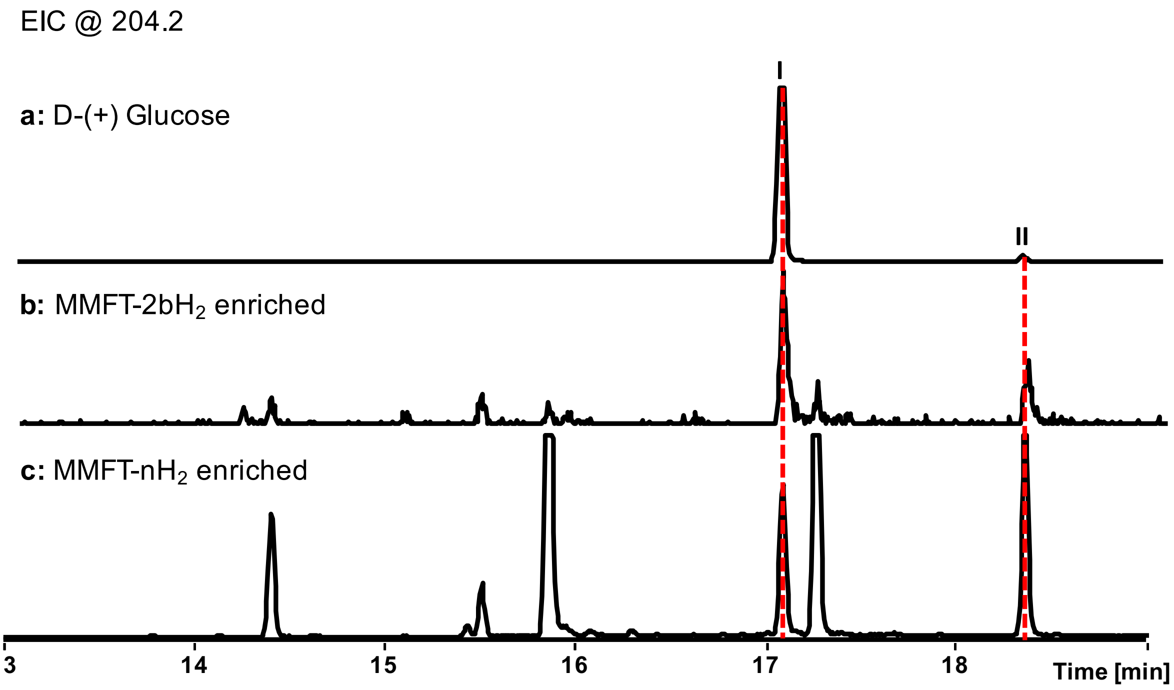


**Supplementary Figure 4. Proposed monosaccharide composition in MMFT.**
**a** GC-MS chromatograms of D-(+)-glucose after MSTFA derivatization. **b** acid hydrolysis of MMFT-2b$H_2$ enriched fractions by 3 M HCl (aq.) and their perTMS derivatives. **c** acid hydrolysis of MMFT-n enriched fractions by 3 M HCl (aq.) and their perTMS derivatives. Compound I ($t_R$ = 17.3 min) and II ($t_R$ = 18.4 min): glucose-1,2,3,4,6-OTMS (GC-MS chromatograms were displayed under EIC mode at *m/z* 204.2, which represented one of the diagnostic fragment ions of hexose-1,2,3,4,6-OTMS. MSTFA: *N*-methyl-*N*-(trimethylsilyl)trifluoroacetamide. TMS: trimethylsilyl.

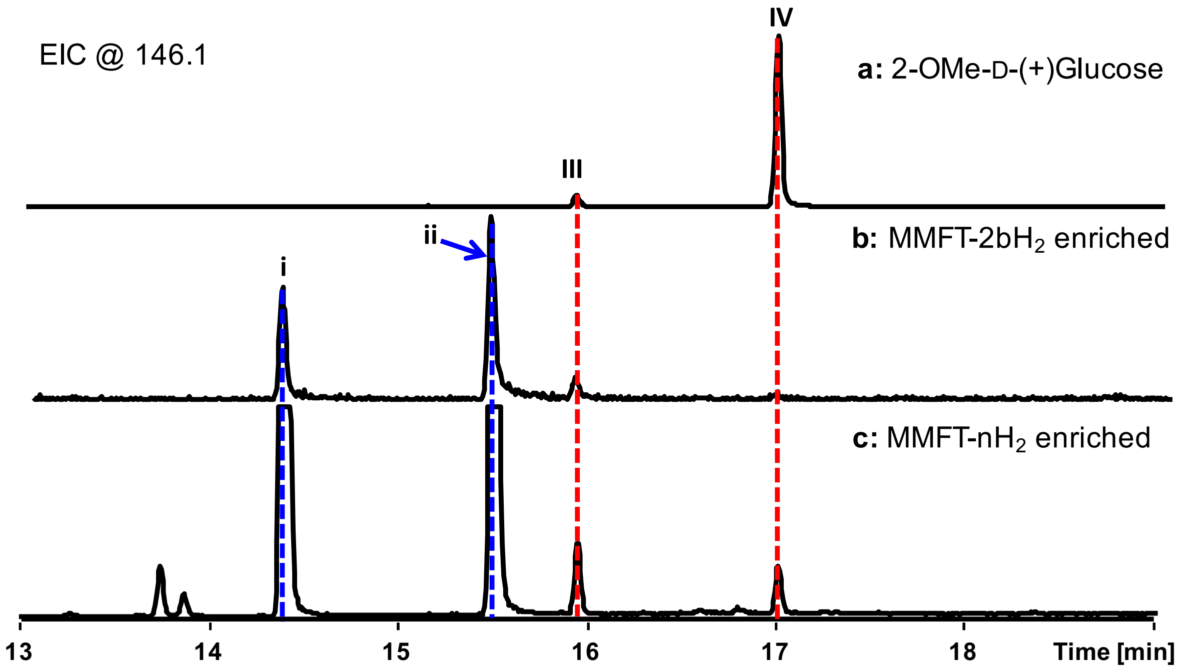


**Supplementary Figure 5. Proposed methylated monosaccharide composition in MMFT.** **a** GC-MS chromatograms of 2-OMe-D-(+)-glucose after MSTFA derivatization. **b** acid hydrolysis of MMFT-2bH$_2$ enriched fractions by 3 M HCl (aq.) and their perTMS derivatives. **c** acid hydrolysis of MMFT-n enriched fractions by 3 M HCl (aq.) and their perTMS derivatives. **Compound III** ($t_R$ = 16.0 min) and **IV** ($t_R$ = 17.0 min): glucose-2-OMe-1,3,4,6-OTMS (GC-MS chromatograms were displayed under EIC mode at *m/z* 146.1, which represented one of the diagnostic fragment ions of hexose-2-OMe-1,3,4,6-OTMS. **Compound i** ($t_R$ = 14.4 min) and **ii** ($t_R$ = 15.5 min): hexose-3-OMe-1,2,4,6-OTMS.

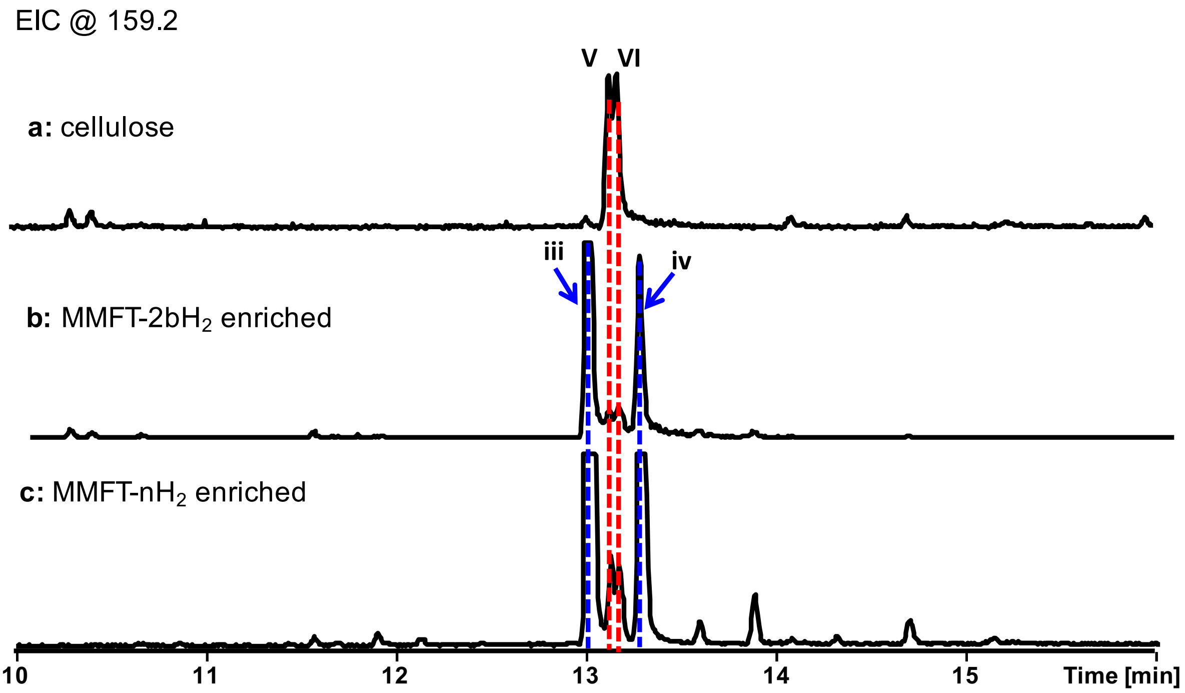


**Supplementary Figure 6. Proposed monosaccharide linkage in MMFT**.
**a** GC-MS chromatogram of cellulose after permethylation and acid hydrolysis by 3 M HCl (aq.) and MSTFA derivatization. **b** MMFT-2b$H_2$ enriched fraction after permethylation and acid hydrolysis by 3 M HCl (aq.) and MSTFA derivatization. **c** MMFT-n enriched fraction after permethylation and acid hydrolysis by 3 M HCl (aq.) and MSTFA derivatization. **Compound V** ($t_R$ = 13.21 min) and **VI** ($t_R$ = 13.26 min) : glucose-2,3,6-OMe-1,4-OTMS (GC-MS chromatograms were displayed under EIC mode at *m*/*z* 159.2, which represented one of the diagnostic fragment ions of hexose-2,3,6-Ome-1, 4-OTMS. **Compound iii** ($t_R$ = 13.0 min) and **iv** ($t_R$ = 13.3 min): hexose-2,3,6-OMe-1,4-OTMS.

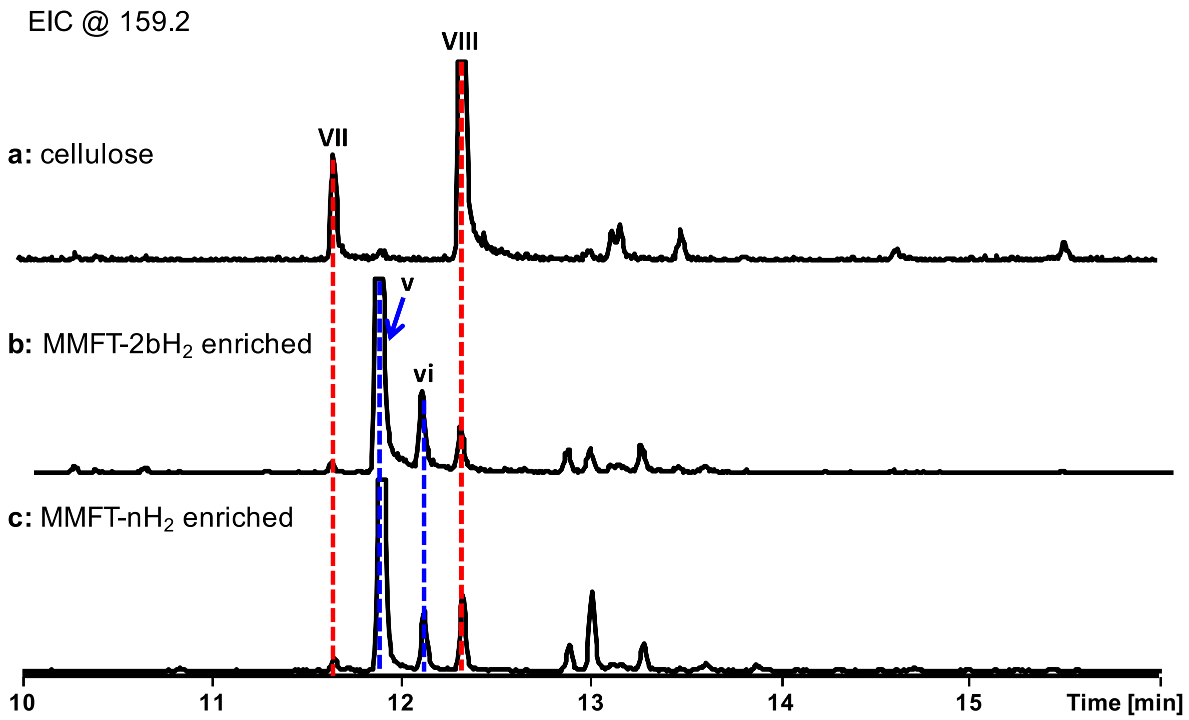


**Supplementary Figure 7. Proposed monosaccharide linkage in MMFT**.
**a** GC-MS chromatogram of cellulose after permethylation and methanolysis by 1.25 M HCl (in MeOH) and MSTFA derivatization. **b** MMFT-2bH$_2$ enriched fraction after permethylation and methanolysis by 1.25 M HCl (in MeOH) and MSTFA derivatization **c** MMFT-n enriched fraction after permethylation and methanolysis by 1.25 M HCl (in MeOH) and MSTFA derivatization. **Compound VII** ($t_R$ = 11.7 min) and **VIII** ($t_R$ = 12.3 min): glucose-1,2,3,6-OMe-4-OTMS (GC-MS chromatograms were displayed under EIC mode at *m*/*z* 159.2, which represented one of the diagnostic fragment ions of hexose-1,2,3,6-OMe-4-OTMS. **Compound v** ($t_R$ = 11.9 min) and **vi** ($t_R$ = 12.1 min): hexose-1,2,3,6-OMe-4-OTMS.

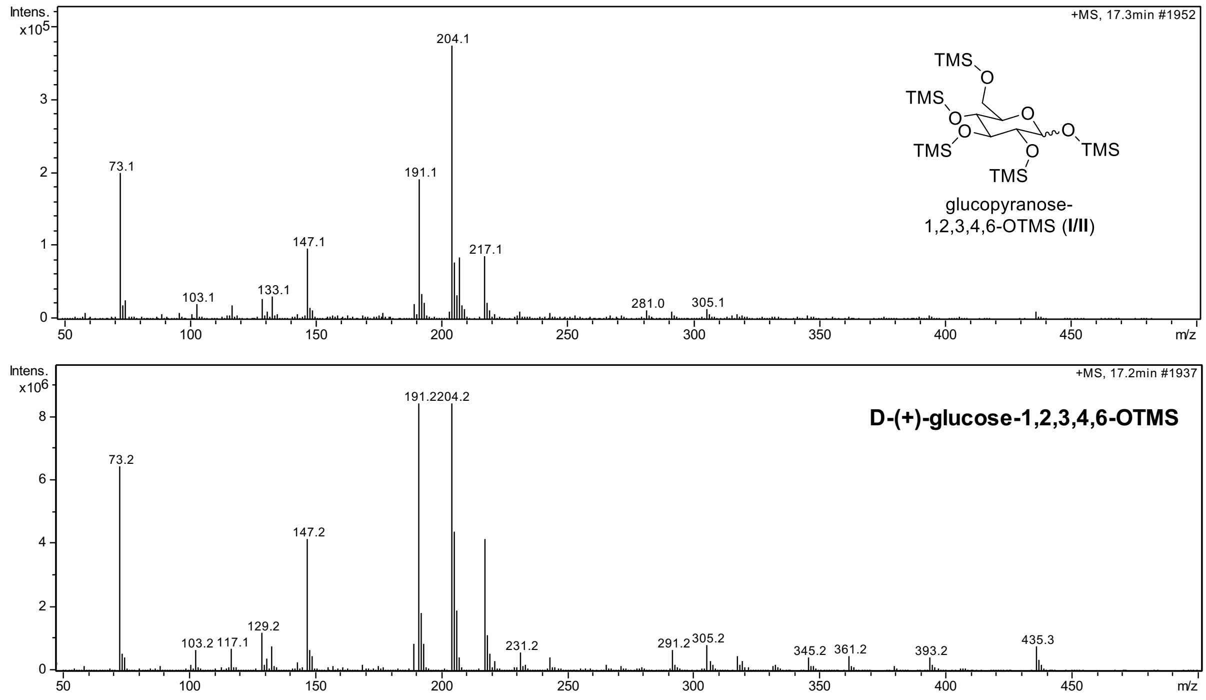


**Supplementary Figure 8**. EI-MS spectra of compound I and standard D-(+)-glucose-1,2,3,4,6-OTMS.

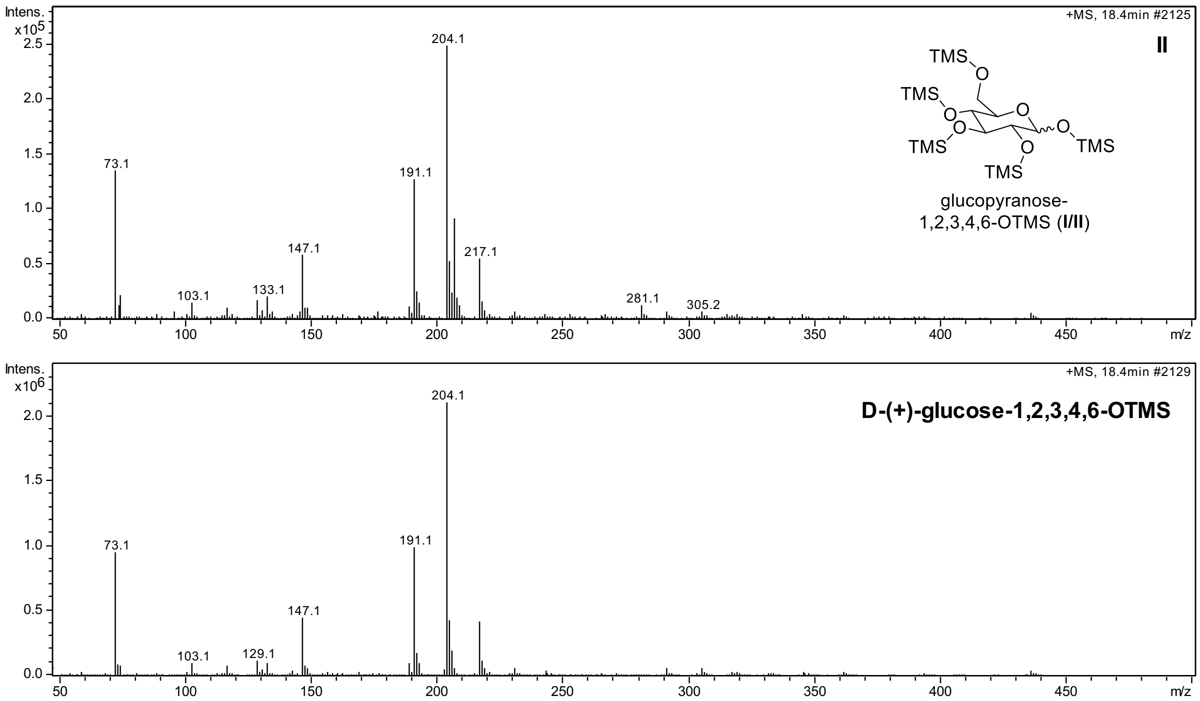

**Supplementary Figure 9**. EI-MS spectra of compound II and standard D-(+)-glucose-1,2,3,4,6-OTMS.

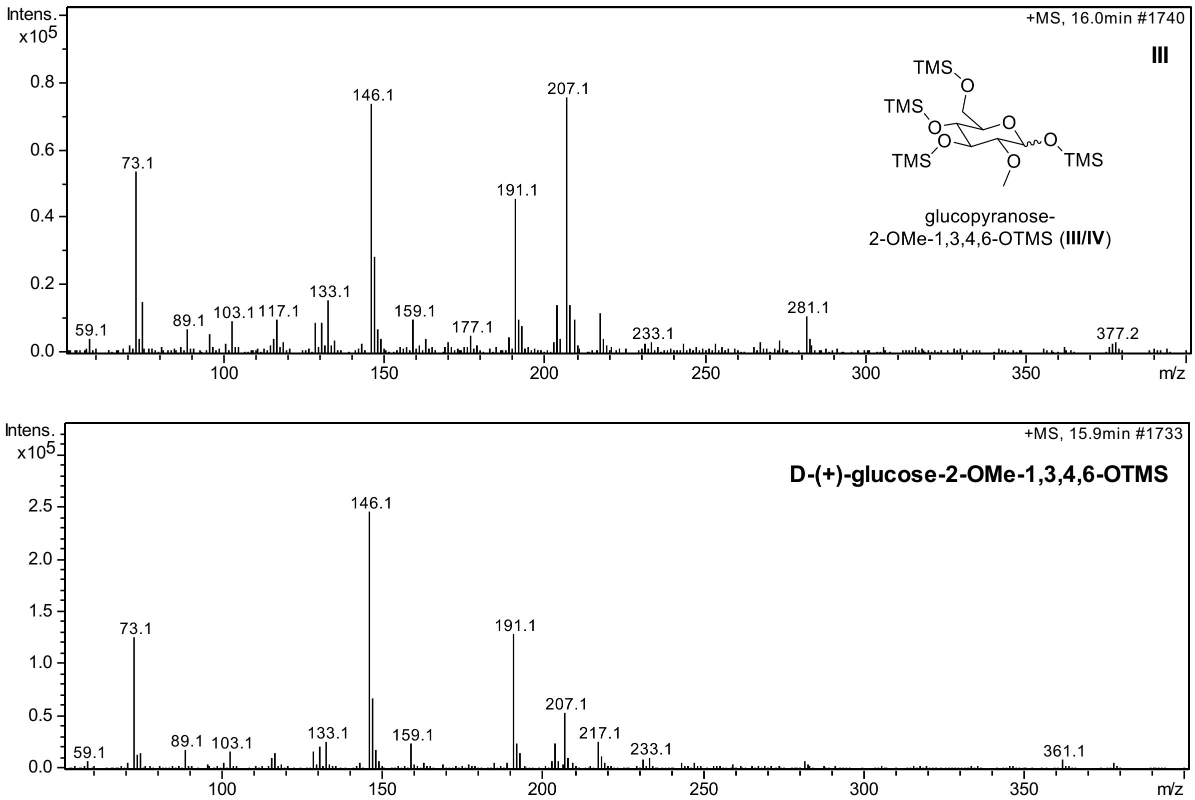


**Supplementary Figure 10.** EI-MS spectra of compound III and standard D-(+)-glucose-2-OMe-1,3,4,6-OTMS

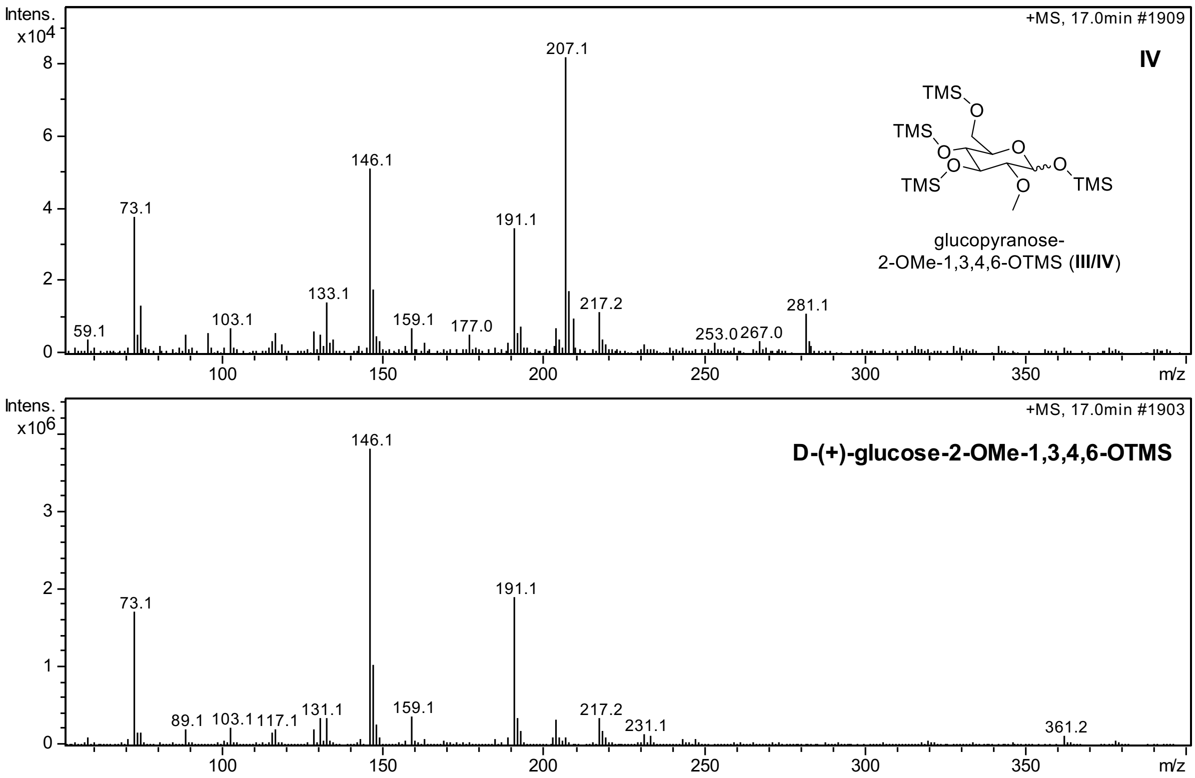


**Supplementary Figure 11**. EI-MS spectra of compound IV and standard D-(+)-glucose-2-OMe-1,3,4,6-OTMS

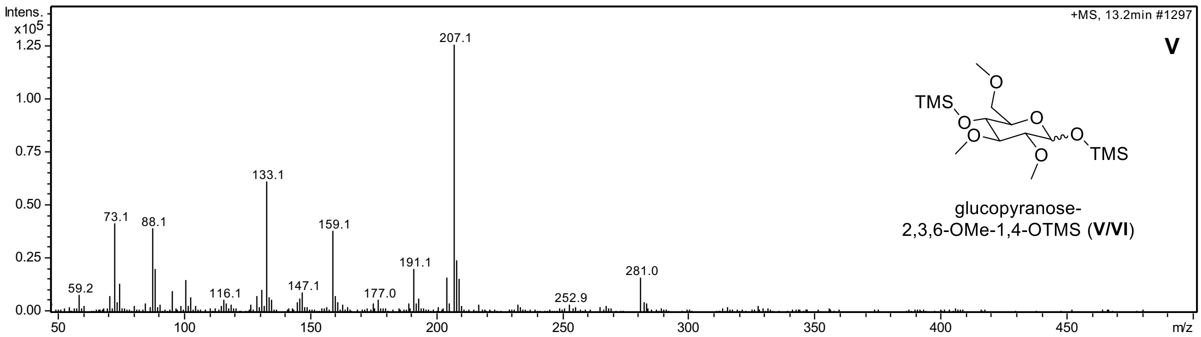


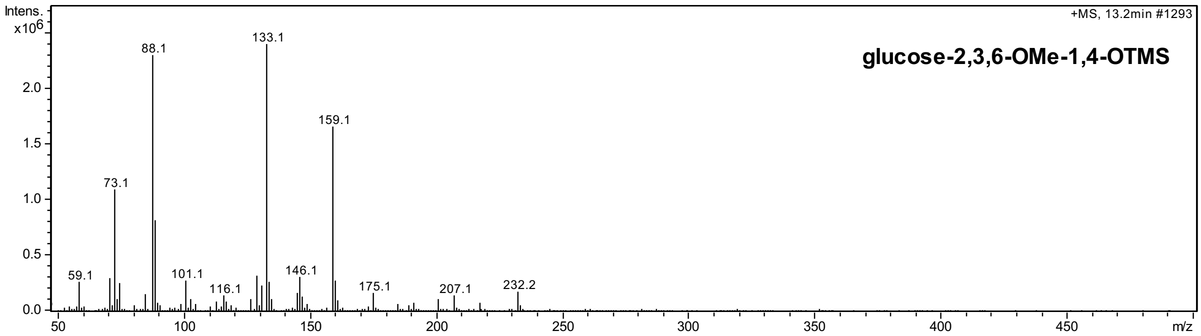


**Supplementary Figure 12**. EI-MS spectra of compound V and glucose-2,3,6-OMe-1,4-OTMS derived from cellulose

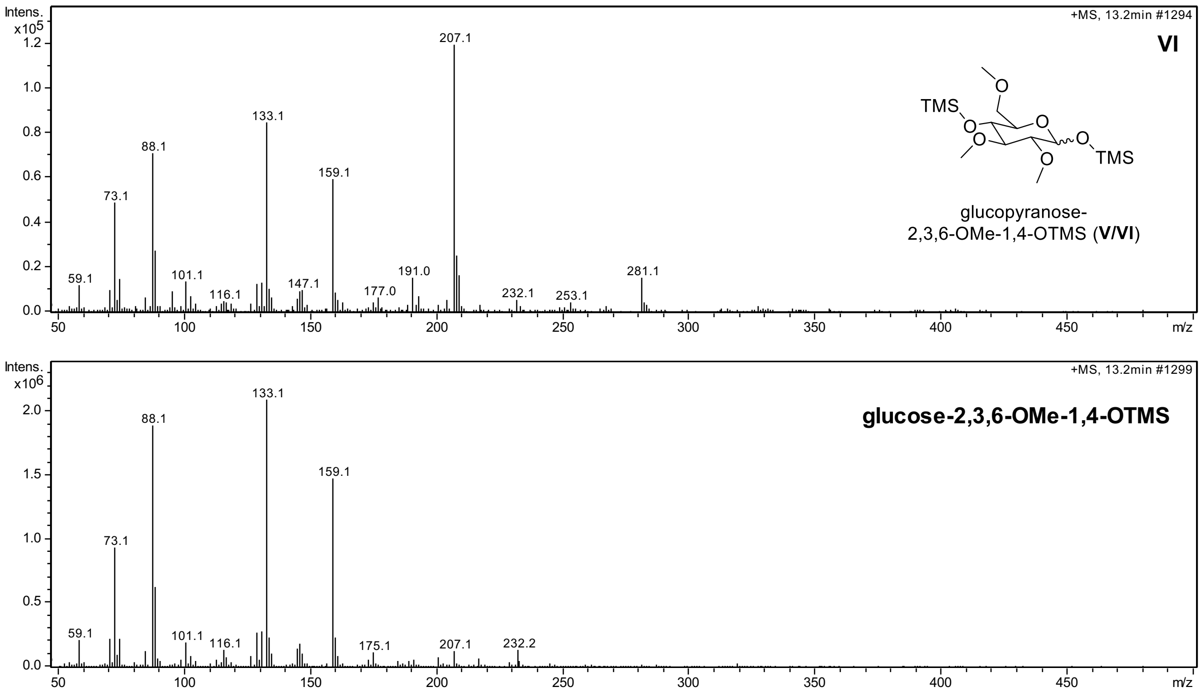


**Supplementary Figure 13**. EI-MS spectra of compound VI and glucose-2,3,6-OMe-1,4-OTMS derived from cellulose

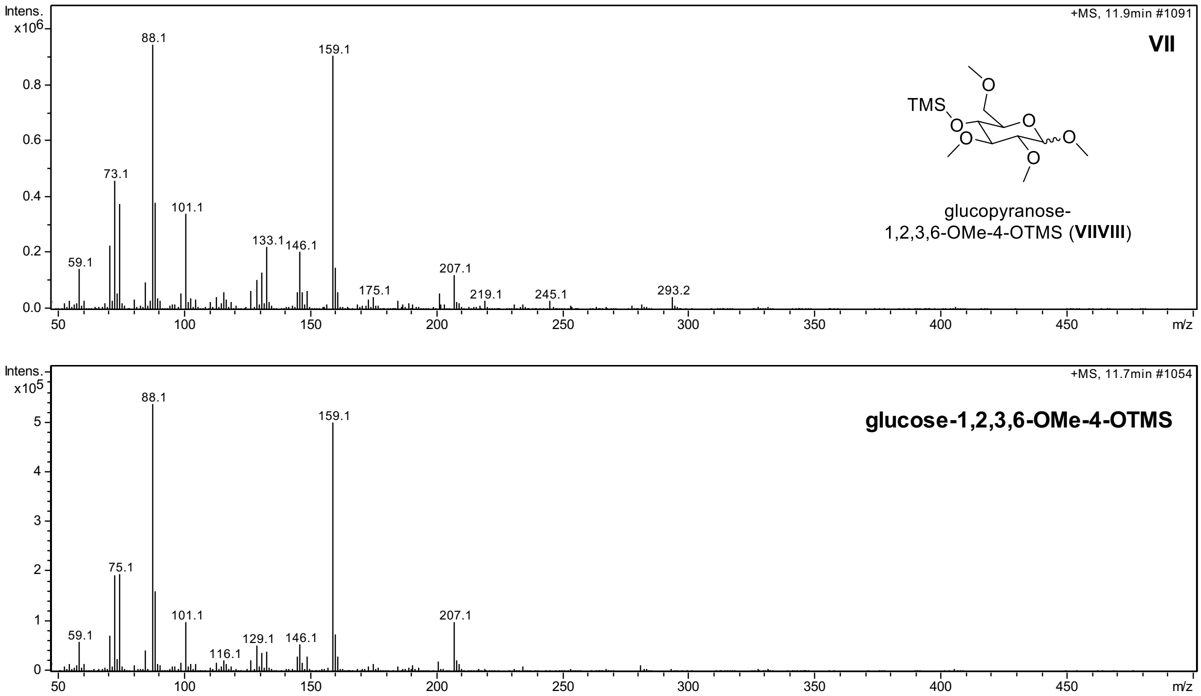


**Supplementary Figure 14**. EI-MS spectra of compound VII and glucose-1,2,3,6-OMe-4-OTMS derived from cellulose

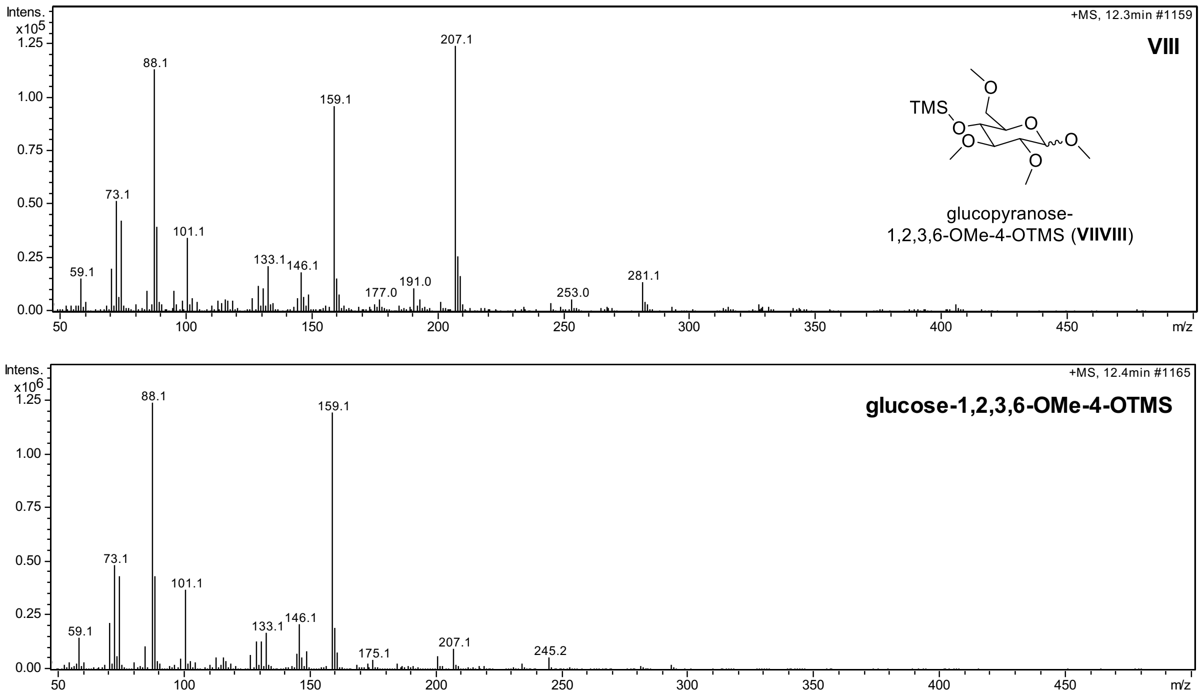


**Supplementary Figure 15**. EI-MS spectra of compound VIII and glucose-1,2,3,6-OMe-4-OTMS derived from cellulose

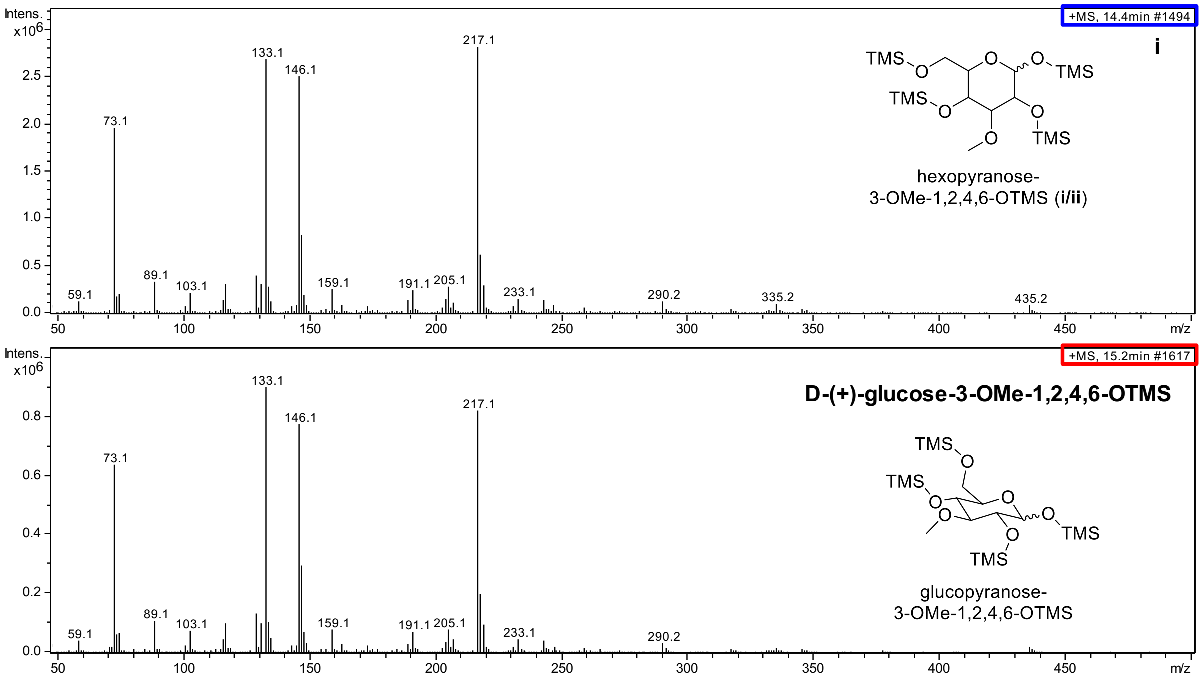


**Supplementary Figure 16**. EI-MS spectra of compound i and standard D-(+)-glucose-3-OMe-1,2,4,6-OTMS

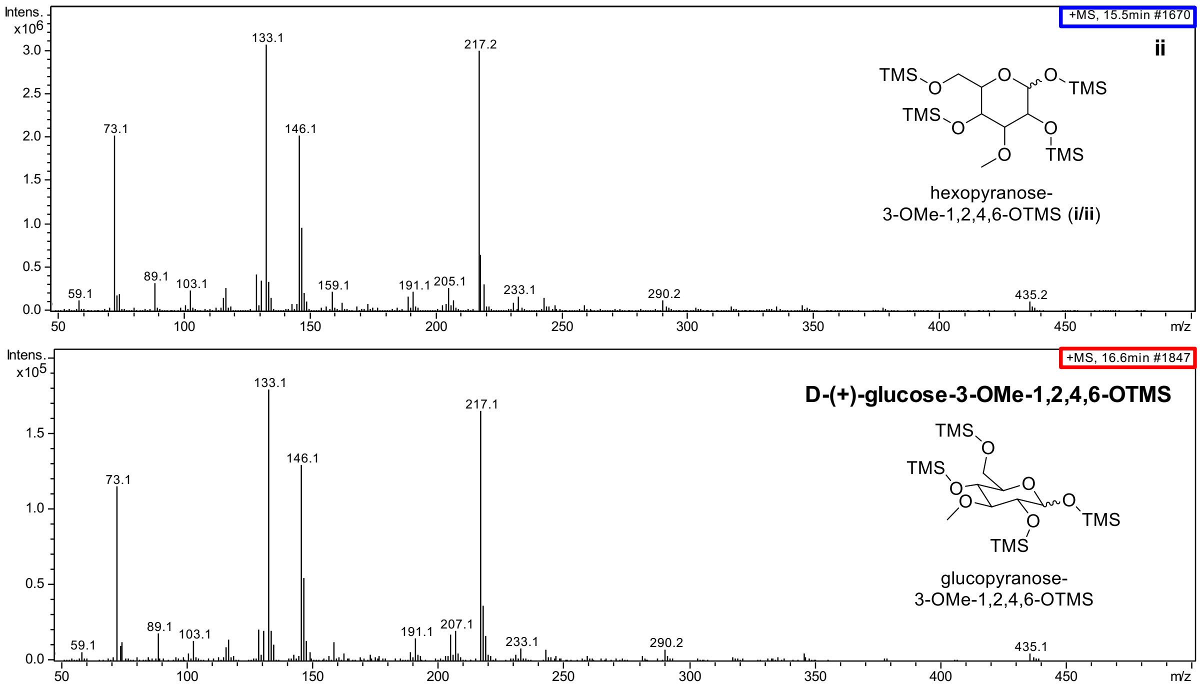


**Supplementary Figure 17**. EI-MS spectra of compound ii and standard D-(+)-glucose-3-OMe-1,2,4,6-OTMS

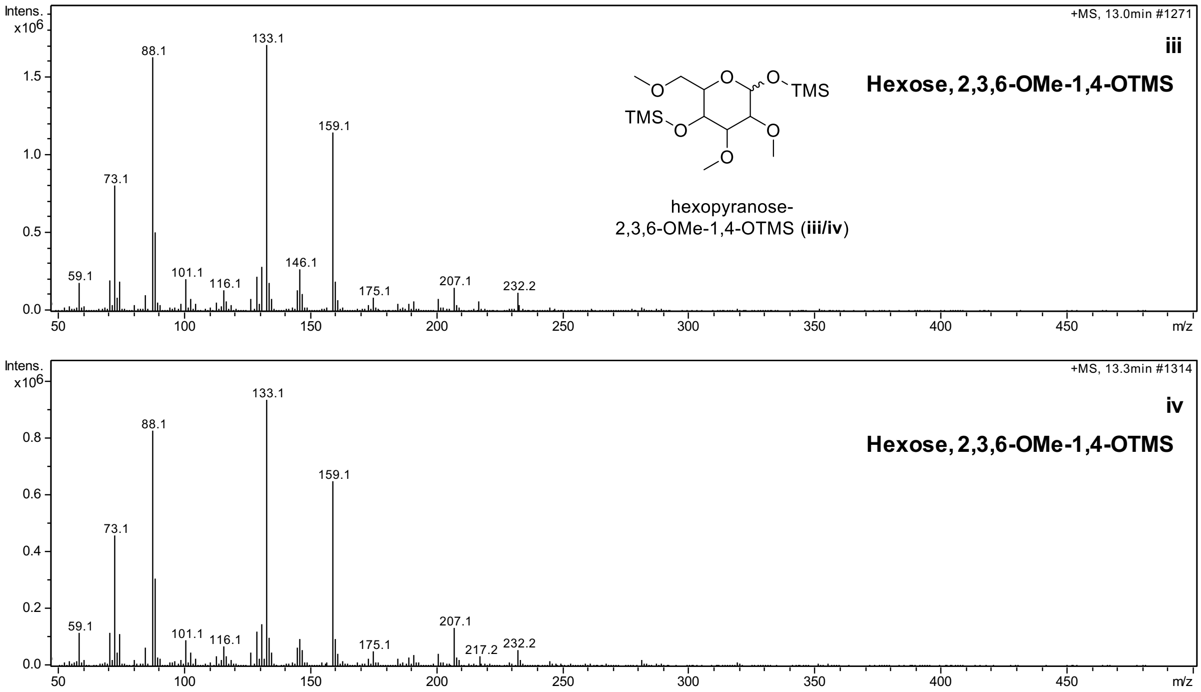


**Supplementary Figure 18**. EIMS spectra of compound iii and iv

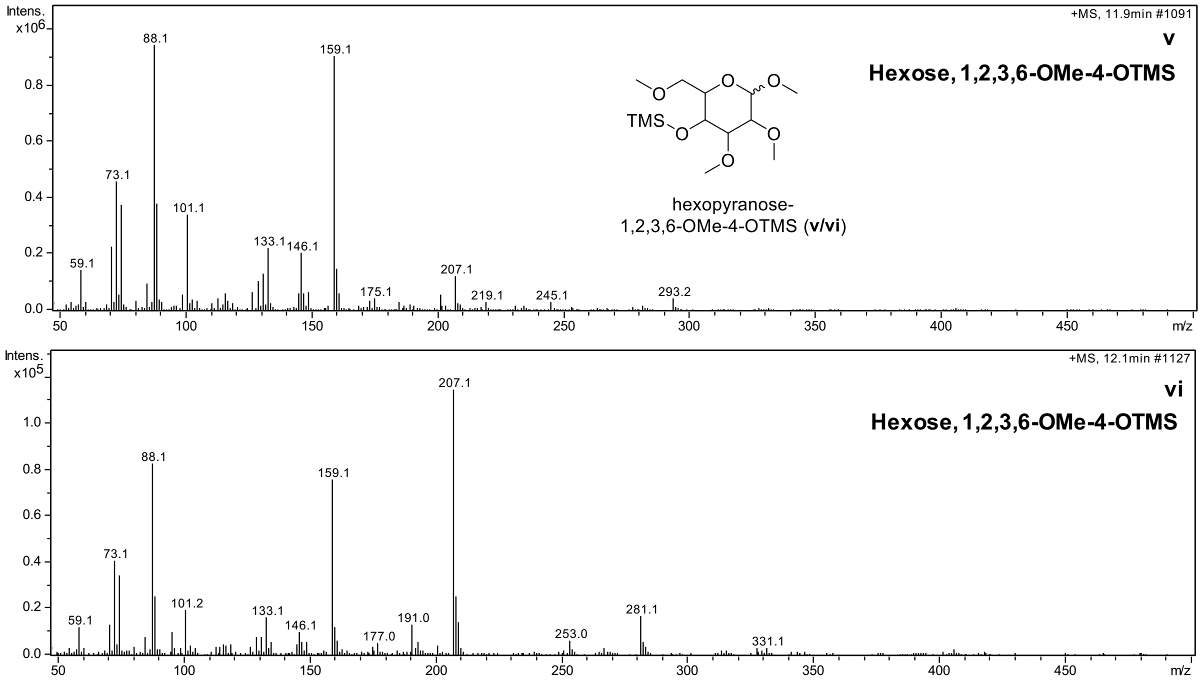


**Supplementary Figure 19**. EI-MS spectra of compound v and vi

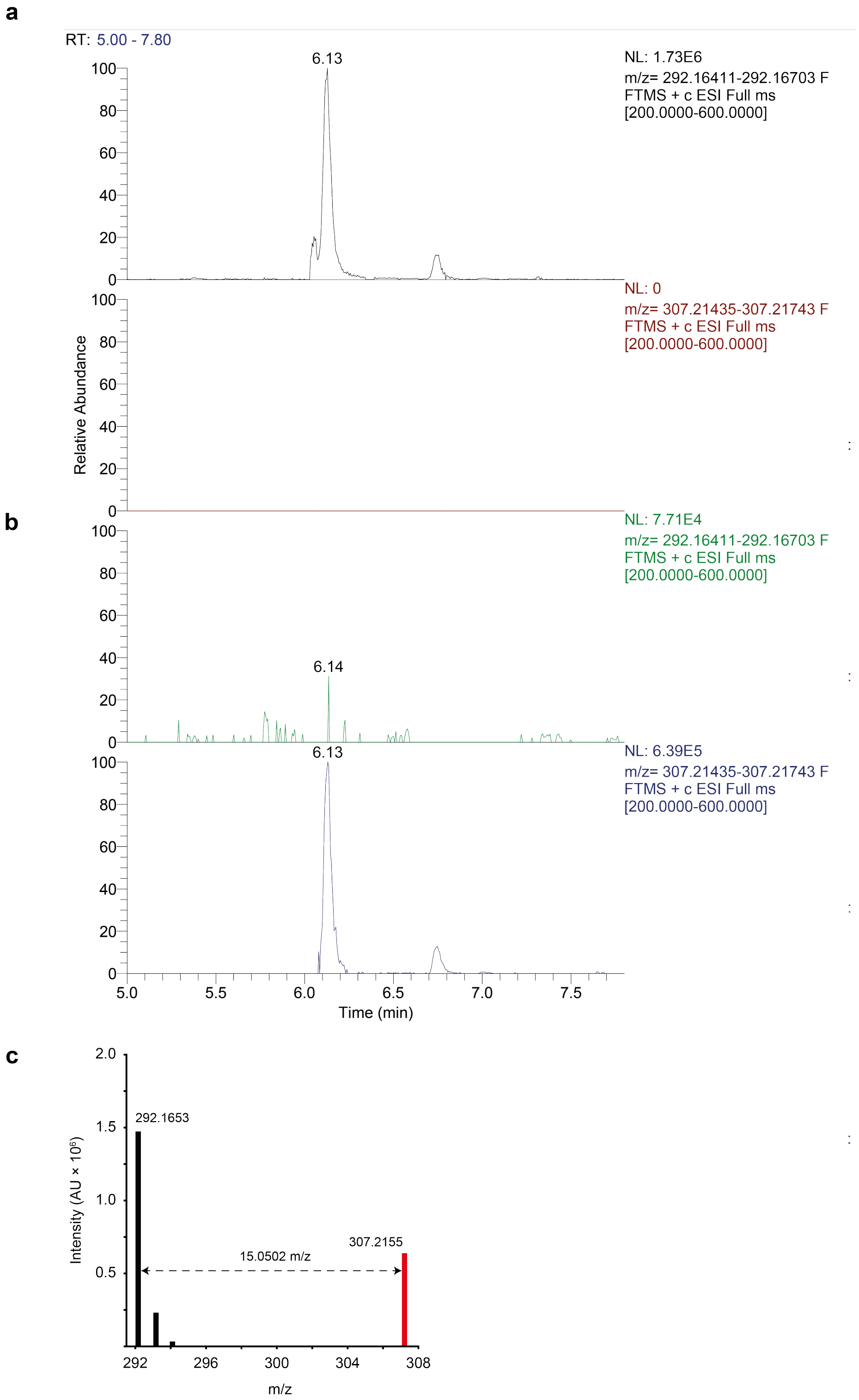


**Supplementary Figure 20. Stable isotope labeling of GAHDP.**
Extracted ion chromatograms of unlabeled (*m/z* 292.16557 $[M+H]^+$) and labeled (*m/z* 307.21589 $[M+H]^+$) GAHDP from **a** *M. smegmatis* fed with unlabeled Gly, Val, and Tyr. **b** *M. smegmatis* fed with $^{13}$C-labeled Gly-$^{13}C_2$, Val-$^{13}C_5$, and Tyr-$^{13}C_9$. **c** Overlaid mass spectrum of unlabeled (black) and labeled GAHDP (red). Labeled GAHDP ($C_{15}H_{21}N_3O_3$) showed a mass shift of +15.05033 Da indicating the incorporation of all three labeled amino acids.

**a**

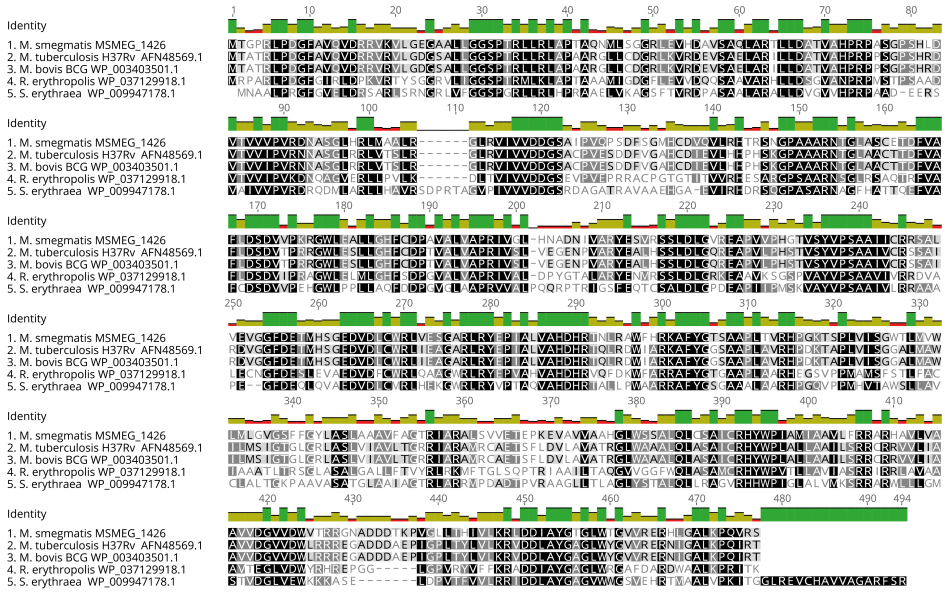


**b**

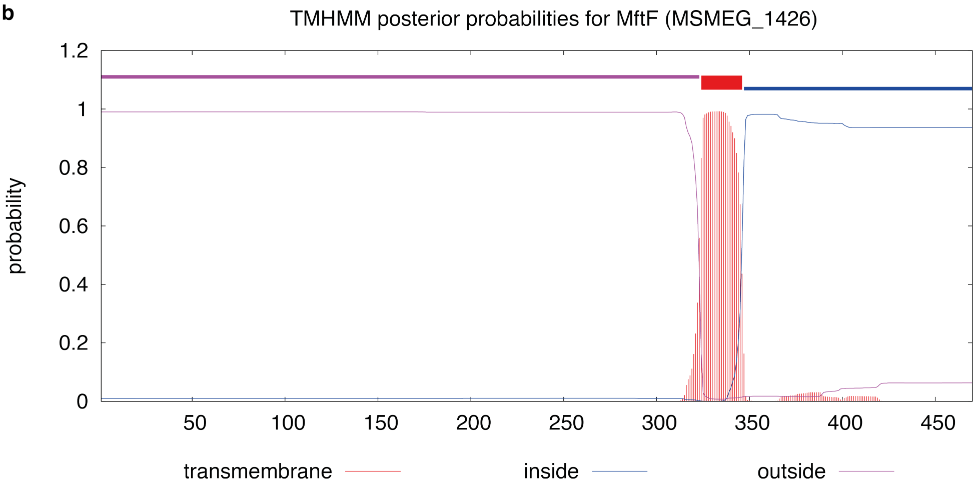


**Supplementary Figure 21**. **Bioinformatics analyses of the MftF protein (MSMEG_1426).**
**a** Sequence alignment (MUSCLE algorithm) of MftF homologues. Source organisms: *Mycolicibacterium smegmatis* MC$^2$ 155, *Mycobacterium tuberculosis* H37Rv, *Mycobacterium bovis* BCG BCG Pasteur 1173P2, *Rhodococcus erythropolis* SK121, *Saccharopolyspora erythraea* NRRL2338. Protein accession numbers are indicated together with sequence names. **b** Transmembrane helix prediction (TMHMM) of MSMEG_1426 revealed a membrane helix spanning residues 24 – 46.

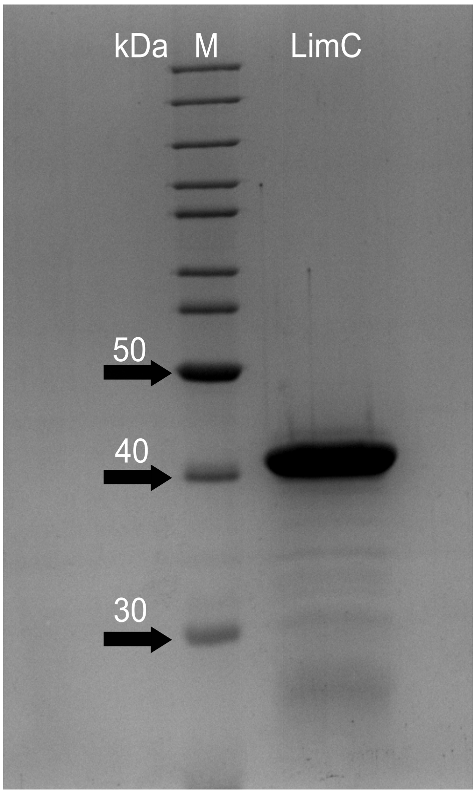


**Supplementary Figure 22. SDS-PAGE of recombinant carveol dehydrogenase LimC from *Rhodococcus erythropolis* DCL14.** The hexahistidin-tagged protein (His6-LimC, 33 kDa) was produced in *E. coli* BL21 (DE3) / pLAPO04 (Supplementary Fig. 23) and purified by nickel-affinity chromatography. The protein elutes at around 40 kDa due to reduced gel mobility. M = molecular weight marker (PageRuler, Thermo Fisher).

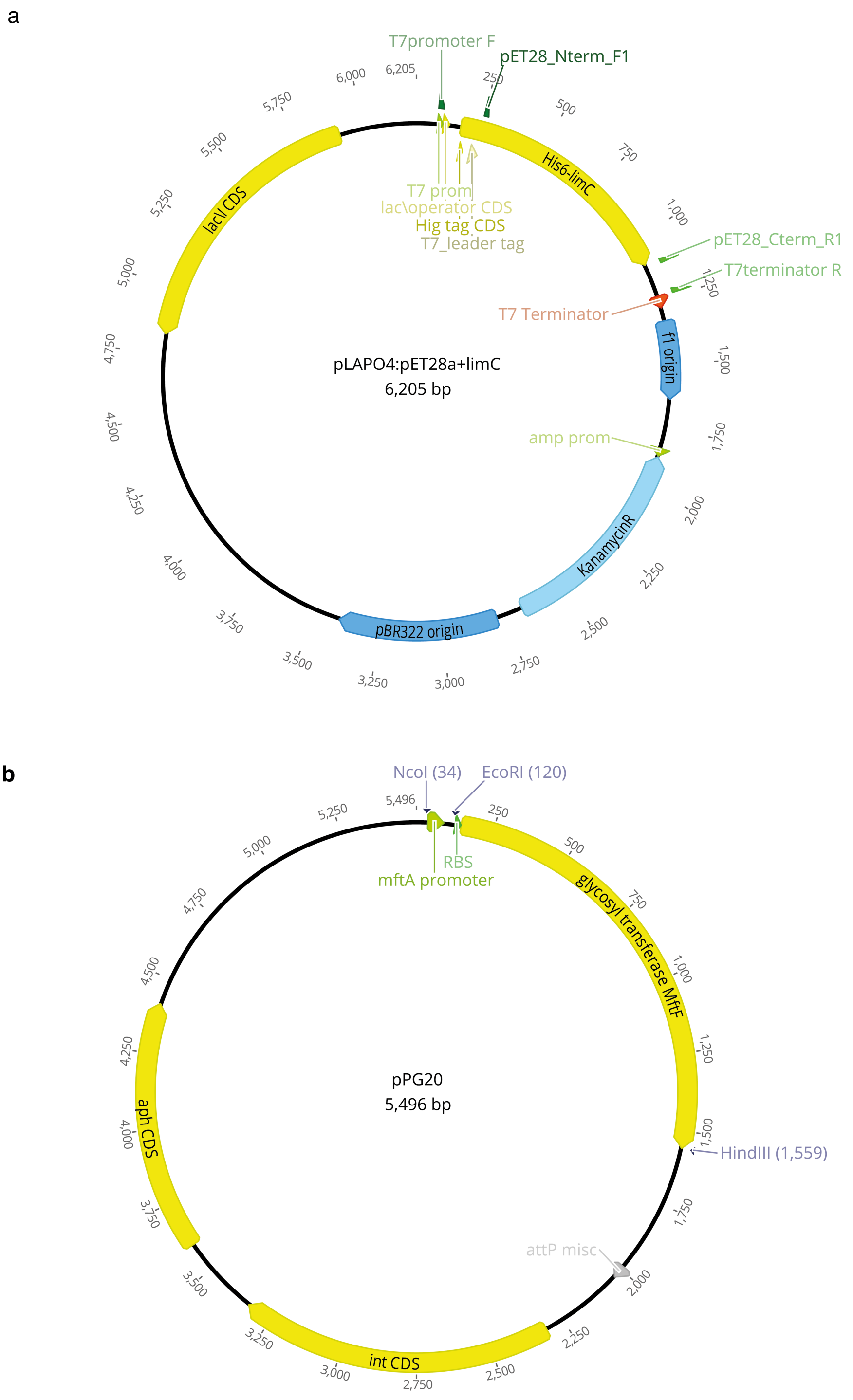


**Supplementary Figure 23. Maps of plasmids used in this study.**
**a** Plasmid pLAPO04 used for heterologous expression of *limC* in *E. coli* BL21 (DE3): Vector pET28a bearing the coding sequence (CDS) of *limC* from *Rhodococcus erythropolis* DCL14 fused to the N-terminal His-tag region. **b** Plasmid pPG20 used for complementation of the *mftF* gene (Δ*mftF*-Comp) in *M. smegmatis*: Integrative vector pMCPAINT carries the promotor region and ribosome binding site (RBS) of *mftA* fused to the natural CDS of *mftF* for homologous expression in *M. smegmatis*. Promotor and RBS are separated by an additional *Eco*RI site. The whole insert was obtained as a synthetic construct flanked by *Nco*I and *Hind*III sites. Plasmid sequences are shown in Section 3.

### 3 Plasmid sequences

**>pLAPO4:pET28a+limC**

GCGCCAGCAACCGCACCTGTGGCGCCGGTGATGCCGGCCACGATGCGTCCGGCGTAGAGGATCGAG
ATCTCGATCCCGCGAAATTAATACGACTCACTATAGGGGAATTGTGAGCGGATAACAATTCCCCTCTA
GAAATAATTTTGTTTAACTTTAAGAAGGAGATATACCATGGGCAGCAGCCATCATCATCATCATCACA
GCAGCGGCCTGGTGCCGCGCGGCAGCCATATGGCTAGCATGACTGGTGGACAGCAAATGGGTCGC
GTCATGGCACGTGTTGAAGGTCAGGTTGCACTGATTACCGGTGCAGCACGTGGTCAGGGTCGTAGC
CATGCAATTAAACTGGCCGAAGAGGGTGCAGATGTTATTCTGGTTGATGTTCCGAATGATGTGGTGG
ATATTGGTTATCCGCTGGGCACCGCAGATGAACTGGATCAGACCGCAAAAGATGTTGAAAATCTGG
GTCGTAAAGCCATTGTTATTCATGCCGATGTTCGTGATCTGGAAAGCCTGACAGCCGAAGTTGATCG
TGCAGTTAGCACCCTGGGTCGTCTGGATATTGTTAGCGCAAATGCAGGTATTGCAAGCGTTCCGTTTC
TGAGCCATGATATTCCGGATAATACCTGGCGTCAGATGATTGATATTAATCTGACCGGTGTTTGGCAT
ACCGCCAAAGTTGCAGTTCCGCATATTCTGGCAGGCGAACGTGGTGGTAGCATTGTTCTGACCAGCA
GCGCAGCAGGTCTGAAAGGTTATGCACAGATTAGCCATTATAGCGCAGCAAAACATGGTGTTGTTG
GTCTGATGCGTAGCCTGGCACTGGAACTGGCACCGCATCGTGTTCGTGTTAATAGCCTGCATCCGAC
ACAGGTTAATACCCCGATGATTCAGAATGAAGGCACCTATCGTATTTTTAGTCCGGACCTGGAAAATC
CGACACGTGAAGATTTTGAAATTGCAAGCACCACCACCAATGCACTGCCGATTCCGTGGGTTGAAAG
CGTTGATGTTAGCAATGCCCTGCTGTTTCTGGTTAGCGAAGATGCACGTTATATTACAGGTGCCGCAA
TTCCGGTTGATGCAGGTACAACCCTGAAATAAGATCCGAATTCGAGCTCCGTCGACAAGCTTGCGGC
CGCACTCGAGCACCACCACCACCACCACTGAGATCCGGCTGCTAACAAAGCCCGAAAGGAAGCTGA
GTTGGCTGCTGCCACCGCTGAGCAATAACTAGCATAACCCCTTGGGGCCTCTAAACGGGTCTTGAGG
GGTTTTTTGCTGAAAGGAGGAACTATATCCGGATTGGCGAATGGGACGCGCCCTGTAGCGGCGCAT
TAAGCGCGGCGGGTGTGGTGGTTACGCGCAGCGTGACCGCTACACTTGCCAGCGCCCTAGCGCCCG
CTCCTTTCGCTTTCTTCCCTTCCTTTCTCGCCACGTTCGCCGGCTTTCCCCGTCAAGCTCTAAATCGGGG
GCTCCCTTTAGGGTTCCGATTTAGTGCTTTACGGCACCTCGACCCCAAAAAACTTGATTAGGGTGATG
GTTCACGTAGTGGGCCATCGCCCTGATAGACGGTTTTTCGCCCTTTGACGTTGGAGTCCACGTTCTTT
AATAGTGGACTCTTGTTCCAAACTGGAACAACACTCAACCCTATCTCGGTCTATTCTTTTGATTTATAA
GGGATTTTGCCGATTTCGGCCTATTGGTTAAAAAATGAGCTGATTTAACAAAAATTTAACGCGAATTT
TAACAAAATATTAACGTTTACAATTTCAGGTGGCACTTTTCGGGGAAATGTGCGCGGAACCCCTATTT
GTTTATTTTTCTAAATACATTCAAATATGTATCCGCTCATGAATTAATTCTTAGAAAAACTCATCGAGC
ATCAAATGAAACTGCAATTTATTCATATCAGGATTATCAATACCATATTTTTGAAAAAGCCGTTTCTGT
AATGAAGGAGAAAACTCACCGAGGCAGTTCCATAGGATGGCAAGATCCTGGTATCGGTCTGCGATT
CCGACTCGTCCAACATCAATACAACCTATTAATTTCCCCTCGTCAAAAATAAGGTTATCAAGTGAGAA
ATCACCATGAGTGACGACTGAATCCGGTGAGAATGGCAAAAGTTTATGCATTTCTTTCCAGACTTGTT
CAACAGGCCAGCCATTACGCTCGTCATCAAAATCACTCGCATCAACCAAACCGTTATTCATTCGTGAT
TGCGCCTGAGCGAGACGAAATACGCGATCGCTGTTAAAAGGACAATTACAAACAGGAATCGAATGC
AACCGGCGCAGGAACACTGCCAGCGCATCAACAATATTTTCACCTGAATCAGGATATTCTTCTAATAC
CTGGAATGCTGTTTTCCCGGGGATCGCAGTGGTGAGTAACCATGCATCATCAGGAGTACGGATAAAA
TGCTTGATGGTCGGAAGAGGCATAAATTCCGTCAGCCAGTTTAGTCTGACCATCTCATCTGTAACATC
ATTGGCAACGCTACCTTTGCCATGTTTCAGAAACAACTCTGGCGCATCGGGCTTCCCATACAATCGAT
AGATTGTCGCACCTGATTGCCCGACATTATCGCGAGCCCATTTATACCCATATAAATCAGCATCCATG
TTGGAATTTAATCGCGGCCTAGAGCAAGACGTTTCCCGTTGAATATGGCTCATAACACCCCTTGTATT
ACTGTTTATGTAAGCAGACAGTTTTATTGTTCATGACCAAAATCCCTTAACGTGAGTTTTCGTTCCACT
GAGCGTCAGACCCCGTAGAAAAGATCAAAGGATCTTCTTGAGATCCTTTTTTTCTGCGCGTAATCTGC
TGCTTGCAAACAAAAAAACCACCGCTACCAGCGGTGGTTTGTTTGCCGGATCAAGAGCTACCAACTC
TTTTTCCGAAGGTAACTGGCTTCAGCAGAGCGCAGATACCAAATACTGTCCTTCTAGTGTAGCCGTAG
TTAGGCCACCACTTCAAGAACTCTGTAGCACCGCCTACATACCTCGCTCTGCTAATCCTGTTACCAGTG
GCT

GCTGCCAGTGGCGATAAGTCGTGTCTTACCGGGTTGGACTCAAGACGATAGTTACCGGATAAGGCG
CAGCGGTCGGGCTGAACGGGGGGTTCGTGCACACAGCCCAGCTTGGAGCGAACGACCTACACCGAA
CTGAGATACCTACAGCGTGAGCTATGAGAAAGCGCCACGCTTCCCGAAGGGAGAAAGGCGGACAG
GTATCCGGTAAGCGGCAGGGTCGGAACAGGAGAGCGCACGAGGGAGCTTCCAGGGGGAAACGCCT
GGTATCTTTATAGTCCTGTCGGGTTTCGCCACCTCTGACTTGAGCGTCGATTTTTGTGATGCTCGTCAG
GGGGGCGGAGCCTATGGAAAAACGCCAGCAACGCGGCCTTTTTACGGTTCCTGGCCTTTTGCTGGCC
TTTTGCTCACATGTTCTTTCCTGCGTTATCCCCTGATTCTGTGGATAACCGTATTACCGCCTTTGAGTG
AGCTGATACCGCTCGCCGCAGCCGAACGACCGAGCGCAGCGAGTCAGTGAGCGAGGAAGCGGAAG
AGCGCCTGATGCGGTATTTTCTCCTTACGCATCTGTGCGGTATTTCACACCGCATATATGGTGCACTCT
CAGTACAATCTGCTCTGATGCCGCATAGTTAAGCCAGTATACACTCCGCTATCGCTACGTGACTGGGT
CATGGCTGCGCCCCGACACCCGCCAACACCCGCTGACGCGCCCTGACGGGCTTGTCTGCTCCCGGCA
TCCGCTTACAGACAAGCTGTGACCGTCTCCGGGAGCTGCATGTGTCAGAGGTTTTCACCGTCATCACC
GAAACGCGCGAGGCAGCTGCGGTAAAGCTCATCAGCGTGGTCGTGAAGCGATTCACAGATGTCTGC
CTGTTCATCCGCGTCCAGCTCGTTGAGTTTCTCCAGAAGCGTTAATGTCTGGCTTCTGATAAAGCGGG
CCATGTTAAGGGCGGTTTTTTCCTGTTTGGTCACTGATGCCTCCGTGTAAGGGGGATTTCTGTTCATG
GGGGTAATGATACCGATGAAACGAGAGAGGATGCTCACGATACGGGTTACTGATGATGAACATGCC
CGGTTACTGGAACGTTGTGAGGGTAAACAACTGGCGGTATGGATGCGGCGGGACCAGAGAAAAAT
CACTCAGGGTCAATGCCAGCGCTTCGTTAATACAGATGTAGGTGTTCCACAGGGTAGCCAGCAGCAT
CCTGCGATGCAGATCCGGAACATAATGGTGCAGGGCGCTGACTTCCGCGTTTCCAGACTTTACGAAA
CACGGAAACCGAAGACCATTCATGTTGTTGCTCAGGTCGCAGACGTTTTGCAGCAGCAGTCGCTTCA
CGTTCGCTCGCGTATCGGTGATTCATTCTGCTAACCAGTAAGGCAACCCCGCCAGCCTAGCCGGGTCC
TCAACGACAGGAGCACGATCATGCGCACCCGTGGGGCCGCCATGCCGGCGATAATGGCCTGCTTCTC
GCCGAAACGTTTGGTGGCGGGACCAGTGACGAAGGCTTGAGCGAGGGCGTGCAAGATTCCGAATA
CCGCAAGCGACAGGCCGATCATCGTCGCGCTCCAGCGAAAGCGGTCCTCGCCGAAAATGACCCAGA
GCGCTGCCGGCACCTGTCCTACGAGTTGCATGATAAAGAAGACAGTCATAAGTGCGGCGACGATAG
TCATGCCCCGCGCCCACCGGAAGGAGCTGACTGGGTTGAAGGCTCTCAAGGGCATCGGTCGAGATC
CCGGTGCCTAATGAGTGAGCTAACTTACATTAATTGCGTTGCGCTCACTGCCCGCTTTCCAGTCGGGA
AACCTGTCGTGCCAGCTGCATTAATGAATCGGCCAACGCGCGGGGAGAGGCGGTTTGCGTATTGGG
CGCCAGGGTGGTTTTTCTTTTCACCAGTGAGACGGGCAACAGCTGATTGCCCTTCACCGCCTGGCCCT
GAGAGAGTTGCAGCAAGCGGTCCACGCTGGTTTGCCCCAGCAGGCGAAAATCCTGTTTGATGGTGG
TTAACGGCGGGATATAACATGAGCTGTCTTCGGTATCGTCGTATCCCACTACCGAGATATCCGCACCA
ACGCGCAGCCCGGACTCGGTAATGGCGCGCATTGCGCCCAGCGCCATCTGATCGTTGGCAACCAGC
ATCGCAGTGGGAACGATGCCCTCATTCAGCATTTGCATGGTTTGTTGAAAACCGGACATGGCACTCC
AGTCGCCTTCCCGTTCCGCTATCGGCTGAATTTGATTGCGAGTGAGATATTTATGCCAGCCAGCCAGA
CGCAGACGCGCCGAGACAGAACTTAATGGGCCCGCTAACAGCGCGATTTGCTGGTGACCCAATGCG
ACCAGATGCTCCACGCCCAGTCGCGTACCGTCTTCATGGGAGAAAATAATACTGTTGATGGGTGTCT
GGTCAGAGACATCAAGAAATAACGCCGGAACATTAGTGCAGGCAGCTTCCACAGCAATGGCATCCT
GGTCATCCAGCGGATAGTTAATGATCAGCCCACTGACGCGTTGCGCGAGAAGATTGTGCACCGCCGC
TTTACAGGCTTCGACGCCGCTTCGTTCTACCATCGACACCACCACGCTGGCACCCAGTTGATCGGCGC
GAGATTTAATCGCCGCGACAATTTGCGACGGCGCGTGCAGGGCCAGACTGGAGGTGGCAACGCCAA
TCAGCAACGACTGTTTGCCCGCCAGTTGTTGTGCCACGCGGTTGGGAATGTAATTCAGCTCCGCCATC
GCCGCTTCCACTTTTTCCCGCGTTTTCGCAGAAACGTGGCTGGCCTGGTTCACCACGCGGGAAACGG
TCTGATAAGAGACACCGGCATACTCTGCGACATCGTATAACGTTACTGGTTTCACATTCACCACCCTG
AATTGACTCTCTTCCGGGCGCTATCATGCCATACCGCGAAAGGTTTTGCGCCATTCGATGGTGTCCGG
GATCTCGACGCTCTCCCTTATGCGACTCCTGCATTAGGAAGCAGCCCAGTAGTAGGTTGAGGCCGTT
GAGCACCGCCGCCGCAAGGAATGGTGCATGCAAGGAGATGGCGCCCAACAGTCCCCCGGCCACGG
GGCCTGCCACCATACCCACGCCGAAACAAGCGCTCATGAGCCCGAAGTGGCGAGCCCGATCTTCCCC
ATCGGTGATGTCGGCGATATAG

**>pPG20**
GGCCGCGGTACCAGATCTTTAAATCTAGATATCCATGGCTCTCACACCCCCTCTTCCATTCTGGCACTC
GATGCCATATATTTGCGATCTCGATCACAACTGTCGAGACCATACGCGAGAATTCGACCGGAACGGA
TTGCTGACATGACCGGACCGAGACTGCCCGACGGTTTTGCCGTGCAGGTGGATCGCCGAGTCAAGG
TGCTCGGAGAGGGCGCGGCGTTGCTGGGCGGCTCACCGACCCGCCTGCTGCGGTTGGCGCCGACG
GCCCAGAACATGCTCAGCGGGGGCCGCCTCGAGGTTCACGACGCGGTGTCCGCGCAACTCGCGCGA
ACCCTCCTGGACGCGACCGTCGCGCATCCGCGGCCTGCCAGCGGGCCGTCCCATCTGGACGTCACAG
TGGTTGTCCCCGTACGGGACAACGCATCCGGCCTGCACCGCCTGATGGCGGCGCTGCGCGGGCTGC
GCGTGATCGTGGTCGACGACGGCTCGGCGATCCCGGTGCAACCGTCCGACTTCTCCGGTATGCACTG
CGACGTGCAGGTGCTGCGGCACACCCGCAGCAACGGCCCCGCGGCCGCGCGTAACACCGGCCTTGC
GTCGTGCGAGACGGATTTCGTGGCGTTCCTCGATTCCGACGTGGTGCCCAAGCGTGGTTGGCTCGAG
GCGCTGCTCGGGCATTTCTGCGATCCGGCCGTGGCACTGGTGGCGCCCCGCATCGTGGGTCTGCACA
ACGCCGACAACATCGTGGCCCGGTACGAGTCCGTCCGATCCTCGCTGGACCTCGGGGTGCGCGAGG
CGCCCGTGGTGCCGCACGGCACGGTGTCGTATGTGCCGAGCGCGGCGATCATCTGCAGGCGGTCGG
CGCTCGTGGAAGTGGGCGGGTTCGACGAGACCATGCACTCGGGGGAGGATGTCGACCTGTGCTGG
CGCCTGGTCGAGTCGGGCGCGCGTCTGCGTTACGAACCGATCGCCCTGGTGGCGCACGATCACCGC
ACCAACCTGCGAGCGTGGTTCCACCGCAAGGCATTCTATGGAACGTCGGCCGCTCCGCTGACGGTGC
GCCATCCCGGTAAGACGTCGCCGCTGGTGATCTCGGGGTGGACGCTGATGGTGTGGCTGATGCTCG
GGGTGGGCTCGTTCTTCGGCTACCTCGCCTCGCTCGCGGCGGCGGTGTTCGCGGGTACCCGCATCGC
GCGGGCGCTCAGCGTCGTCGAGACCGAACCCAAGGAGGTCGCGGTGGTCGCCGCCCACGGCCTGTG
GTCGTCGGCGTTGCAGTTGTGTTCGGCGATCTGTCGCCACTACTGGCCCATCGCGATGATCGCGGCG
GTGCTGTTCCGCCGGGCGCGGCACGCGGTGCTGGTGGCCGCGGTGGTGGACGGTGTGGTCGACTG
GGTGACGCGACGCGGCAACGCCGACGACGACACCAAACCGGTCGGACTGCTCACCCACATCGTGCT
CAAGCGACTGGACGACATCGCCTACGGCACCGGTCTGTGGACCGGCGTGGTGCGCGAGCGTCACCT
CGGCGCGCTCAAGCCCCAGGTGCGGAGTTAGAAGCTTATCGATGTCGACGTAGTTAACTAGCGTAC
GATCGACTGCCAGGCATCAAATAAAACGAAAGGCTCAGTCGAAAGACTGGGCCTTTCGTTTTATCTG
TTGTTTGTCCGGCCATCATGGCCGCGGTGATCAGCTAGAGCCGTGAACGACAGGGCGAACGCCAGC
CCGCCGACGGCGAGGGTTCCGACCGCTGCAACTCCCGGTGCAACCTTGTCCCGGTCTATTCTCTTCAC
TGCACCAGCTCCAATCTGGTGTGAATGCCCCTCGTCTGTTCGCGCAGGCGGGGGGCTCTATTCGTTT
GTCAGCATCGAAAGTAGCCAGATCAGGGATGCGTTGCAACCGCGTATGCCCAGGTCAGAAGAGTCG
CACAAGAGTTGCAGACCCCTGGAAAGAAAAATGGCCAGAGGGCGAAAACACCCTCTGACCAGCGGA
GCGGGCGACGGGAATCGAACCCGCGTAGCTAGTTTGGAAGAATGGGTGTCTGCCGACCACATATAT
GGGCCGGTCAAGATAGGTTTTTACCCCCTCTCGGCTGCATCCTCTAAGTGGAAAGAAATTGCAGGTC
GTAGAAGCGCGTTGAAGCCTGAGAGTTGCACAGGAGTTGCAACCCGGTAGCCTTGTTCACGACGAG
AGGAGACCTAGTTGGCACGTCGCGGATGGGGATCGCTGAAGACTCAGCGCAGCGGGAGGATCCAA
GCCTCATACGTCAACCCGCAGGACGGTGTGAGGTACTACGCGCTGCAGACCTACGACAACAAGATG
GACGCCGAAGCCTGGCTCGCGGGCGAGAAGCGGCTCATCGAGATGGAGACCTGGACCCCTCCACAG
GACCGGGCGAAGAAGGCAGCCGCCAGCGCCATCACGCTGGAGGAGTACACCCGGAAGTGGCTCGT
GGAGCGCGACCTCGCAGACGGCACCAGGGATCTGTACAGCGGGCACGCGGAGCGCCGCATCTACCC
GGTGCTAGGTGAAGTGGCGGTCACAGAGATGACGCCAGCTCTGGTGCGTGCGTGGTGGGCCGGGA
TGGGTAGGAAGCACCCGACTGCCCGCCGGCATGCCTACAACGTCCTCCGGGCGGTGATGAACACAG
CGGTCGAGGACAAGCTGATCGCAGAGAACCCGTGCCGGATCGAGCAGAAGGCAGCCGATGAGCGC
GACGTAGAGGCGCTGACGCCTGAGGAGCTGGACATCGTCGCCGCTGAGATCTTCGAGCACTACCGG
ATCGCGGCATACATCCTGGCGTGGACGAGCCTCCGGTTCGGAGAGCTGATCGAGCTTCGCCGCAAG
GACATCGTGGACGACGGCATGACGATGAAGCTCCGGGTGCGCCGTGGCGCTTCCCGCGTGGGGAAC
AAGATCGTCGTTGGCAACGCCAAGACCGTCCGGTCGAAGCGTCCTGTGACGGTTCCGCCTCACGTCG
CGGAGATGATCCGAGCGCACATGAAGGACCGTACGAAGATGAACAAGGGCCCCGAGGCATTCCTG
GTGACCACGACGCAGGGCAACCGGCTGTCGAAGTCCGCGTTCACCAAGTCGCTGAAGCGTGGCTAC

GCCAAGATCGGTCGGCCGGAACTCCGCATCCACGACCTCCGCGCTGTCGGCGCTACGTTCGCCGCTC
AGGCAGGTGCGACGACCAAGGAGCTGATGGCCCGTCTCGGTCACACGACTCCTAGGATGGCGATGA
AGTACCAGATGGCGTCTGAGGCCCGCGACGAGGCTATCGCTGAGGCGATGTCCAAGCTGGCCAAGA
CCTCCTGAAACGCAAAAAGCCCCCCTCCCAAGGACACTGAGTCCTAAAGAGGGGGGTTTCTTGTCAG
TACGCGAAGAACCACGCCTGGCCGCGAGCGCCAGCACCGCCGCTCTGTGCGGAGACCTGGGCACCA
GCCCCGCCGCCGCCAGGAGCATTGCCGTTCCCGCCAGCTAGCAACAAAGCGACGTTGTGTCTCAAAA
TCTCTGATGTTACATTGCACAAGATAAAAATATATCATCATGAACAATAAAACTGTCTGCTTACATAA
ACAGTAATACAAGGGGTGTTATGAGCCATATTCAACGGGAAACGTCTTGCTCGAGGCCGCGATTAAA
TTCCAACATGGATGCTGATTTATATGGGTATAAATGGGCTCGCGATAATGTCGGGCAATCAGGTGCG
ACAATCTATCGCTTGTATGGGAAGCCCCATGCGCCAGAGTTGTTTCTGAAACATGGCAAAGGTAGCG
TTGCCAATGATGTTACAGATGAGATGGTCAGACTAAACTGGCTGACGGAATTTATGCCTCTTCCGACC
ATCAAGCATTTTATCCGTACTCCTGATGATGCATGGTTACTCACCACTGCGATCCCCGGGAAAACAGC
ATTCCAGGTATTAGAAGAATATCCTGATTCAGGTGAAAATATTGTTGATGCGCTGGCAGTGTTCCTGC
GCCGGTTGCATTCGATTCCTGTTTGTAATTGTCCTTTTAACAGCGATCGCGTATTTCGTCTCGCTCAGG
CGCAATCACGAATGAATAACGGTTTGGTTGATGCGAGTGATTTTGATGACGAGCGTAATGGCTGGCC
TGTTGAACAAGTCTGGAAAGAAATGCATAATCTTTTGCCATTCTCACCGGATTCAGTCGTCACTCATG
GTGATTTCTCACTTGATAACCTTATTTTTGACGAGGGGAAATTAATAGGTTGTATTGATGTTGGACGA
GTCGGAATCGCAGACCGATACCAGGATCTTGCCATCCTATGGAACTGCCTCGGTGAGTTTTCTCCTTC
ATTACAGAAACGGCTTTTTCAAAAATATGGTATTGATAATCCTGATATGAATAAATTGCAGTTTCATTT
GATGCTCGATGAGTTTTTCTAATCAGAATTGGTTAATTGGTTGTAACACTGGCAGAGCATTACGCTGA
CTTGACGGGACGGCGGCTTTGTTGAATAAATCGAACTTTTGCTGAGTTGAAGGATCAGATCACGCAT
CTTCCCGACAACGCAGACCGTTCCGTGGCAAAGCAAAAGTTCAAAATCACCAACTGGTCCACCTACA
ACAAAGCTCTCATCAACCGTGGCTCCCTCACTTTCTGGCTGGATGATGGGGCGATTCAGGCCTGGTA
TGAGTCAGCAACACCTTCTTCACGAGGCAGACCTCACTAGTTCCACTGAGCGTCAGACCCCGTAGAA
AAGATCAAAGGATCTTCTTGAGATCCTTTTTTTCTGCGCGTAATCTGCTGCTTGCAAACAAAAAAACC
ACCGCTACCAGCGGTGGTTTGTTTGCCGGATCAAGAGCTACCAACTCTTTTTCCGAAGGTAACTGGCT
TCAGCAGAGCGCAGATACCAAATACTGTCCTTCTAGTGTAGCCGTAGTTAGGCCACCACTTCAAGAA
CTCTGTAGCACCGCCTACATACCTCGCTCTGCTAATCCTGTTACCAGTGGCTGCTGCCAGTGGCGATA
AGTCGTGTCTTACCGGGTTGGACTCAAGACGATAGTTACCGGATAAGGCGCAGCGGTCGGGCTGAA
CGGGGGGTTCGTGCACACAGCCCAGCTTGGAGCGAACGACCTACACCGAACTGAGATACCTACAGC
GTGAGCATTGAGAAAGCGCCACGCTTCCCGAAGGGAGAAAGGCGGACAGGTATCCGGTAAGCGGC
AGGGTCGGAACAGGAGAGCGCACGAGGGAGCTTCCAGGGGGAAACGCCTGGTATCTTTATAGTCCT
GTCGGGTTTCGCCACCTCTGACTTGAGCGTCGATTTTTGTGATGCTCGTCAGGGGGGCGGAGCCTAT
GGAAAAACGCCAGCAACGCGGCCTTTTTACGGTTCCTGGCCTTTTGCTGGCCTTTTGCTCACATGTTC
TTTCCTGCGTTATCCCCTGATTCTGTGGATAACCGTATTACCGCCTTTGAGTGAGCTGATACCGCTCGC
CGCAGCCGAACGACCGAGCGCAACGCGTGC
